# Low-pass sequencing increases the power of GWAS and decreases measurement error of polygenic risk scores compared to genotyping arrays

**DOI:** 10.1101/2020.04.29.068452

**Authors:** Jeremiah H. Li, Chase A. Mazur, Tomaz Berisa, Joseph K. Pickrell

## Abstract

Low-pass sequencing (sequencing a genome to an average depth less than 1× coverage) combined with genotype imputation has been proposed as an alternative to genotyping arrays for trait mapping and calculation of polygenic scores. To empirically assess the relative performance of these technologies for different applications, we performed low-pass sequencing (targeting coverage levels of 0.5× and 1×) and array genotyping (using the Illumina Global Screening Array (GSA)) on 120 DNA samples derived from African and European-ancestry individuals that are part of the 1000 Genomes Project. We then imputed both the sequencing data and the genotyping array data to the 1000 Genomes Phase 3 haplotype reference panel using a leave-one-out design. We evaluated overall imputation accuracy from these different assays as well as overall power for GWAS from imputed data, and computed polygenic risk scores for coronary artery disease and breast cancer using previously derived weights. We conclude that low-pass sequencing plus imputation, in addition to providing a substantial increase in statistical power for genome wide association studies, provides increased accuracy for polygenic risk prediction at effective coverages of ~ 0.5× and higher compared to the Illumina GSA.

## Introduction

Thousands of variants on the human genome associated with hundreds of complex traits and diseases have been reproducibly and robustly identified since the first large GWASs for complex disease were performed in the mid-00s (Sella and Barton, 2019; Visscher et al., 2012; Consortium et al., 2007; Yang et al., 2010). Results from these studies have had an enormous impact on the understanding of the genetic architecture underlying complex traits in humans, with these studies playing an essential part in bringing the current understanding full circle back to Fisher’s original infinitesimal model compared to the gene-centric model focused upon in the preceding decades (Walsh and Lynch, 2018; Risch et al., 1999; Judson, 1979).

The ability to systematically dissect the genetic architecture of complex traits influenced by hundreds or thousands of genetic variants has largely been enabled by the dense genotyping array, which cost-effectively assays the genome of an individual at hundreds of thousands to millions of loci (Visscher et al., 2012). Imputation of the resulting genotypes to existing haplotype reference panels further allows evaluation of genetic variants which are not directly assayed, often resulting in total callsets many times the number of directly assayed loci, and is now standard practice in preparing genomic datasets for GWAS (Li et al., 2009; Marchini and Howie, 2010).

As genome sequencing costs have decreased over the past decade, sequencing-based alternatives to genotyping arrays have been the subject of growing interest (Wetterstrand, 2019). Specifically, low-coverage shotgun whole genome sequencing followed by imputation has been utilized for a number of problems in statistical and population genetics, from providing the backbone for graph-based pangenomes in sorghum to trait mapping in human pharmacogenetics (Tran et al., 2020; Gilly et al., 2019; Rubinacci et al., 2020; Homburger et al., 2019; Wasik et al., 2019; Jensen et al., 2020; Cai et al., 2015a; Liu et al., 2018). As an intuition for why this approach is useful, a sample sequenced at a target coverage of 0.5× is expected to have at least one read on 33 million of the 85 million sites in the 1000 Genomes Phase 3 release, whereas a genotyping array will probe a number of variants which is one to two orders of magnitude fewer, albeit with higher average accuracy (Consortium et al., 2015).

For many use cases, there are a number of advantages to low-pass sequencing (lps) over genotyping arrays; for example, (1) there is a lack of ascertainment bias with regard to which variants/sites on the genome are assayed, (2) sequence data can be used to discover novel variation both at the sample or population level (such as in Tran et al. (2020); Liu et al. (2018)), (3) massively parallel low-pass sequencing can be achieved by multiplexing large numbers of samples to reduce cost, and (4) the fact that the average expected accuracy of a sample’s imputed genotypes can be fine-tuned by adjusting the target coverage for the sample, something which is useful when designing experiments within real-world logistical or budgeting constraints (see Pasaniuc et al. (2012) for a detailed simulation-based cost-benefit analysis of low-pass sequencing compared to genotyping arrays for GWAS study designs).

However, studies in the literature which investigate the applications of low-pass sequencing often do so by means of simulation or by *downsampling* (often already-aligned) sequence reads from samples previously sequenced at higher coverages (Gilly et al., 2019; Homburger et al., 2019). While useful, these approaches are unable to capture the real-world idiosyncrasies of data generation in extremely-low coverage sequencing and ignore factors such as the use of different library preparation methods which are optimized for sequencing at given target coverages (Aird et al., 2011; Jones et al., 2015).

Here, in order to more realistically represent real-world results, we perform an investigation of low-pass sequencing data where the low-coverage genomes are obtained not by downsampling higher-coverage samples but by direct sequencing to extremely low target coverages (0.5× and 1.0×). As a point of comparison, we chose to assay the same samples on the Illumina Global Screening Array (GSA), a modern genotyping array specifically designed to capture multi-ethnic genetic variation. We chose these coverages since the all-in cost of library preparation, sequencing, and data analysis for a 0.5× target coverage sample was approximately at parity with the Illumina GSA (as of 2019), and that for 1.0× target coverage sequencing fast approaching the same with historically dropping sequencing costs (Wetterstrand, 2019).

## Results

### Experimental Overview

In order to compare the relative performance of low-pass sequencing and genotyping arrays across populations, we selected 60 EUR and 60 AFR individuals (Supplementary Table 1) from the 1000 Genomes Phase 3 release (1KGP3) (Consortium et al., 2015) on which to perform five experiments (Table 1, Supplementary Table 2), which we denote experiments A-E:

**Table 1:** Details of experiments conducted. Experiments A-D were based on low-pass sequencing (lpSeq) while experiment E used the Illumina GSA v3.0. The samples column describes the number of unique cell lines and the number of replicates run for each of them. Library prep was performed by Gencove for experiments A-C and libraries were sequenced on the Illumina HiSeqX. For experiment D, DNA was sent to BGI who performed the library prep and sequencing. Empirical coverage for each sample was calculated by dividing the number of bases sequenced by the size of the human genome (~ 3.3Gb).

| Experiment | Mean Coverage | Samples | Assay | Library Prep | Sequencer |
| --- | --- | --- | --- | --- | --- |
| A | 0.67 | 120 (3 replicates) | lpSeq | KAPA HyperPlus | Illumina HiSeqX |
| B | 1.25 | 120 (3 replicates) | lpSeq | KAPA HyperPlus | Illumina HiSeqX |
| C | 1.20 | 1 (30 replicates) | lpSeq | KAPA HyperPlus | Illumina HiSeqX |
| D | 1.26 | 60 (1 replicate) | lpSeq | MGIEasy | BGISEQ 500 |
| E (array) | NA | 120 (3 replicates) | Illumina GSA v3.0 | NA | NA |

For experiment A, we performed library prep on and sequenced these 120 unique individuals in triplicate to a target coverage of 0.5× on an Illumina HiSeqX. For experiment B, we performed library prep on and sequenced these 120 unique individuals in triplicate to a target coverage of 1.0× on an Illumina HiSeqX. For experiment C, we performed library prep on and sequenced NA12878, a CEU female sample, thirty times to a target coverage of 1.0× on an Illumina HiSeqX. For experiment D, we selected a subset of 30 EUR and 30 AFR samples from the set of 120 unique individuals and sent DNA to BGI Americas to be sequenced a target coverage of 1.0× on a BGIseq 500. For experiment E, we assayed these 120 unique individuals in triplicate on the Illumina GSA v3.0 via the Broad Institute.

Experiment C was conducted principally to illustrate the effects of varying empirical coverage on imputation accuracy (with all else held equal) and to provide insight into the repeatability of low-pass sequencing on biological replicates and empirical variation in sequencing coverage given a fixed target coverage (Supplementary Figure 1).

For each assayed sample passing QC (Methods, Supplementary Table 3), we imputed the sequence or genotype array data to the 1KGP3 haplotype reference panel in a leave-one-out manner (Methods) and compared the imputed calls against the left-out genotypes from the 1KGP3 reference panel (which we treated as the “gold-standard” or “truth” set). We also computed polygenic risk scores from the imputed dosages for each sample and the gold-standard truth set for breast cancer (BC) and coronary artery disease (CAD) using variant weights from recent state-of-the-art studies (Mavaddat et al., 2019; Inouye et al., 2018).

### Defining “effective coverage”

The nominal (mapped) coverage of a sample having undergone whole genome sequencing is defined as the number of sequenced (and mapped) bases divided by the size of the genome (in this case, ~ 3.3Gb). This quantity is useful as an indicator of how much sequence data is available for downstream analysis, but does not give any useful information as to how spatially uniform (with respect to the genome coodinate system) the sequenced data are distributed. Spatial uniformity of sequencing reads is particularly important for low-pass sequencing followed by imputation because imputation panels catalogue variation across the entire genome and the imputation quality at a given variant is influenced by the amount of sequence data mapped to regions near that variant and which overlap other variants in the imputation panel.

We therefore introduce the concept of a sequenced sample’s *effective coverage λ*_eff_, which is a function of the fraction of polymorphic sites in a haplotype reference panel covered by at least one sequencing read. Under an idealized Poisson distribution of sequencing reads across sites, this fraction is determined by the sequencing coverage alone (Methods). Specifically, given an imputation panel with *n* sites and a set of aligned reads from a single sample, we can compute the fraction of those *n* sites covered by at least one read *f*_covered_ and compute that sample’s effective coverage *λ*_eff_ = − ln(1 − *f*_covered_).

The advantage of using this quantity rather than nominal coverage as a way to summarize sequence results on a sample, particularly at ultra-low coverages, is that the assumptions of an idealised sampling process is “built-in” to its definition. This allows results from, for example, different library preparation methods to be compared on more equal footing (see Figure 1, Supplementary Figures 1, 2, and 3, where experiment D underwent a different library preparation method than experiments A-C).

**Figure 1:**
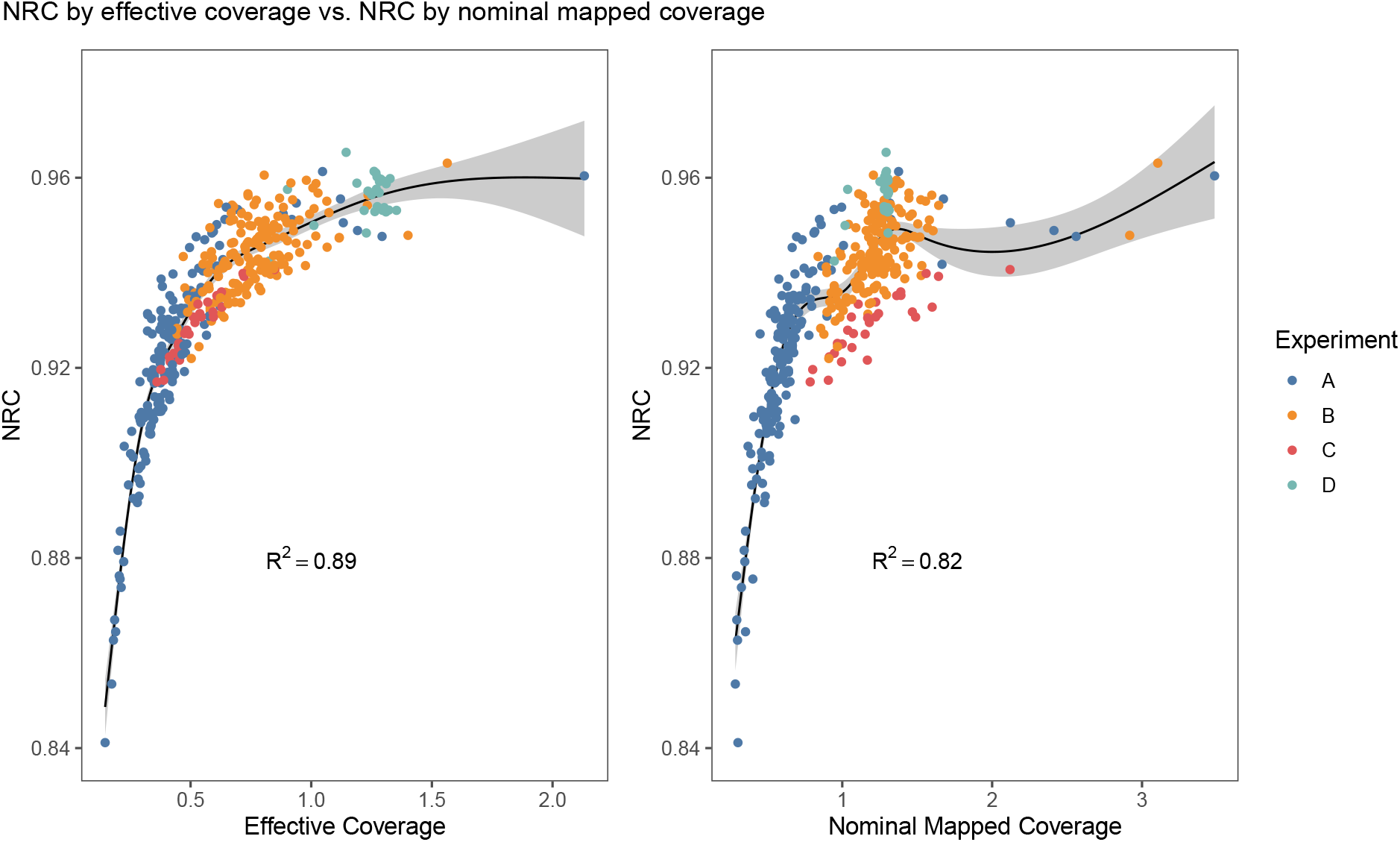
Non-reference concordance (NRC) (Methods) for imputed SNPs for all EUR samples in Experiments A-D plotted against effective coverage *λ*_eff_ (left pane) or nominal mapped coverage (right pane). We modeled the NRC as the response variable for a *k* = 5 knot cubic restricted spline with the respective coverage metric as the explanatory variable; the fitted values are shown as a solid line with the surrounding shaded regions representing 95% confidence intervals. The knot locations were set at the 5th, 27.5th, 50th, 72.5th, and 95th percentiles of the respective coverage metrics following Harrell’s rule of thumb (Harrell, 2017).

Indeed, plotting nominal and effective coverage vs non-reference concordance (NRC) (Methods) for all the sequenced samples (Figure 1) illustrates how NRC is better predicted by a sample’s effective coverage rather than their nominal mapped coverage, with cubic restricted spline fits explaining a larger degree of variance when considering effective coverage rather than mapped nominal coverage (*R*^2^ = 0.89 vs *R*^2^ = 0.82). Note that this figure and accompanying fit are not meant to be a rigorous parametric treatment of NRC vs. different coverage metrics, but are rather meant to provide an intuition.

The results from experiment C illustrate that this pattern also holds across replicates of the same individual, where the only degree of freedom left is the effective coverage of the sample.

Plots of effective coverage vs NRC and overall concordance broken down by variant type (SNPs vs indels), population, and variant filtration status are shown in Supplementary Figures 4, 5, 6, 7, 8, and 9, and show qualitatively similar results. Notably, when poorly imputed variants are filtered out of a callset (variants with a maximum genotype probability of less than 90%, Methods), the relationship between effective coverage and concordance weakens significantly, suggesting that the genotype posterior probabilities generated during imputation are relatively well-calibrated (Methods).

### Comparison of imputation quality metrics across experiments

#### Genotype Concordance

We then examined how NRC and imputation *r*^2^ varied across experiments A, B, D, and E. In order to do this, for experiments A, B, and E, we took a random sample of one of the replicates run for each unique cell line, such that comparisons across experiments concerned only a single, representative sample of each cell line per experiment (Methods). For experiment D we retained all samples as there were no replicates. The remainder of subsection compares metrics and results from these *representative cohorts* across experiments. The mean effective coverage of the representative cohorts varied from experiment to experiment (Supplementary Table 4), ranging from an overall (mean ± standard deviation) of 0.42 ± 0.22 for experiment A to 0.71 ± 0.17 for experiment B to 1.24 ± 0.11 for experiment D.

Reference to non-reference and minor allele frequencies are with respect to those found in the 1KGP3. For all analyses, we treated the genotypes in the 1KGP3 for each sample’s cell line as the gold-standard “truth” set.

We observed that imputed genotypes in EUR cohorts were considerably and consistently more accurate than those imputed into the AFR cohorts on average both before and after filtering out poorly imputed variants both for SNPs and indels, with the difference in mean NRC for unfiltered SNPs ranging from 0.0143 in experiment D to to 0.08 in experiment E (Table 2). Within each experiment, two-sample unpaired Welch’s *t*-tests for differences in means between results for EUR and AFR samples all yielded significant (*P* < 0.001) results.

**Table 2:** Mean and interquartile range (in parentheses) non-reference concordance (NRC) across samples for each representative cohort for unfiltered SNPs by experiment and super population.

| Super Population | Experiment |  |  |  |
| --- | --- | --- | --- | --- |
|  | A | B | D | E (array) |
| AFR | 0.9002 (0.8896-0.9089) | 0.9217 (0.9123-0.9288) | 0.9419 (0.9378-0.9434) | 0.8308 (0.8119-0.8425) |
| EUR | 0.9227 (0.913-0.9342) | 0.9438 (0.9397-0.9477) | 0.9562 (0.9531-0.9594) | 0.9067 (0.898-0.9116) |

Furthermore, we observed that genotypes imputed from sequence were consistently and considerably more accurate than those imputed from array data, with the mean AFR NRC being 7% higher for sequence data with an average of 0.4× effective coverage compared to the array data (Table 2, Supplementary Tables 5, 6, 7). Two-sample paired *t*-tests for differences in means between the array results and sequence results in Table 2 (given a superpopulation) all yielded significant (*P* < 0.001) results.

In order to compare performance across the allele frequency spectrum, we computed the mean non-reference concordance for SNPs within a given allele frequency bin within each representative cohort (Figure 2, Supplementary Figure 10, Supplementary Table 12). The average NRCs for experiment E’s representative cohorts were significantly lower than those of all other experiments at all frequency bins, and qualitatively similar patterns hold for *overall* genotype concordance as well (Supplementary Tables 8, 9, 10, 11). These results indicate that low-pass sequencing at an effective coverage of ~ 0.4× or higher consistently yields more accurate imputed genotype calls at sites of common variation than the Illumina GSA and that this pattern holds across both EUR and AFR cohorts.

**Figure 2:**
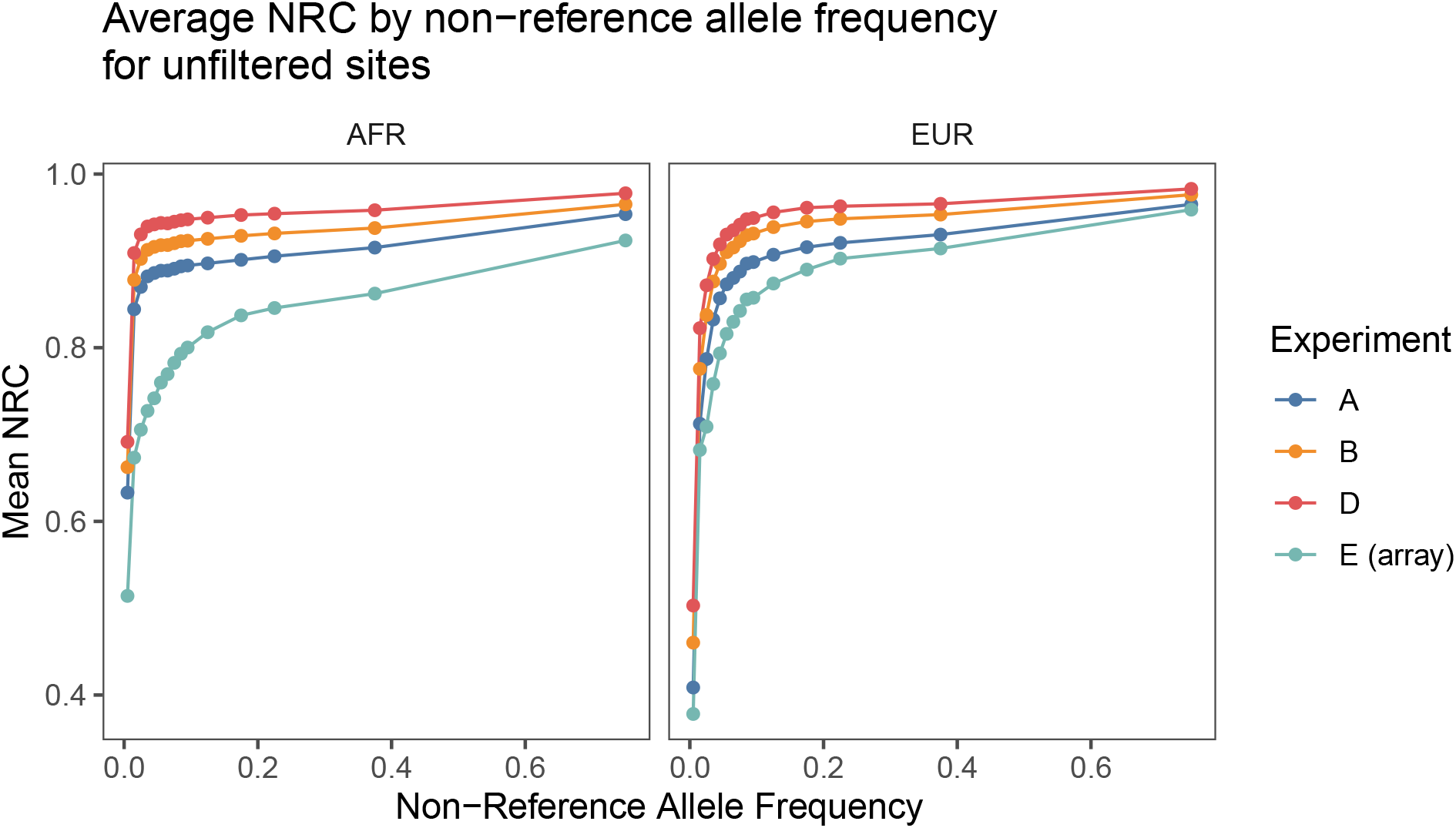
Average non-reference concordance for unfiltered SNPs by superpopulation by non-reference allele frequency in 1KGP3. The NRC for imputed sequence data was consistently higher than the NRC for the imputed GSA data across the allele frequency spectrum. Filtering to confidently imputed variants reveals a similar pattern (Supplementary Figure 10).

In particular, the NRC at low (< 5%) allele frequencies in AFR populations often exceeds the corresponding NRC for EUR populations (Supplementary Figure 11) for the sequence-based experiments. We also observe this pattern for the imputation *r*^2^s, which we address next.

#### Imputation *r*^2^

Genotype concordance is an important metric which provides a straightforward and intuitive quantification of the performance of genotype imputation. However, in the context of genome-wide association studies (GWAS), a more relevant quantity is the *imputation r*^2^, which is defined as the squared correlation between the imputed dosage and the true genotypes of a given set of samples at a given variant. This is because the imputation *r*^2^ at a given variant is proportional to the expectation of the *χ*^2^ statistic resulting from an association test at that particular variant, of which the power of the test is a function (Marchini, 2019; Chapman et al., 2003; Pritchard and Przeworski, 2001). A practical consequence of this is that a proportional increase in the imputation *r*^2^ at a particular variant can be interpreted as the same proportional increase in effective sample size at that variant, so that an increase in mean imputation *r*^2^ across the genome directly corresponds to increased statistical power (Pritchard and Przeworski, 2001). This quantity can be computed on a site-by-site basis and stratified into allele frequency bins according to the allele frequencies in the haplotype reference panel.

At common variants (minor allele frequency > 5% in the 1KGP3), the mean imputation *r*^2^ for sequence-based experiments ranged from 0.92 − 0.96 for the EUR representative cohorts compared to a mean *r*^2^ of 0.91 for experiment E, representing an average increase in power of ~ 1 − 6% for samples with mean effective coverages ranging from ~ 0.47 − 1.24× (Supplementary Tables 4, 15). For the AFR representative cohorts, the mean imputation *r*^2^ for sequence-based experiments ranged from 0.89 − 0.95 compared to 0.83 for the GSA, representing an average increase of power of ~ 7 − 15% for samples with mean effective coverages ranging from ~ 0.38 − 1.24 (Supplementary Table 15). For all experiments and superpopulations at common variants, the mean dosage *r*^2^ for the imputed sequence was significantly higher (*P* < 0.001, two-sample Welch’s *t*-tests) than the corresponding mean dosage *r*^2^ for the imputed array data.

Stratifying imputation site-wise *r*^2^s into minor allele frequency bins, we observed that for the sequence-based experiments, the average *r*^2^s across the allele frequency bins scaled with the average effective coverage of each representative cohort (Figure 3), in line with the expectation that higher-effective-coverage sequence data affords greater imputation accuracy.

**Figure 3:**
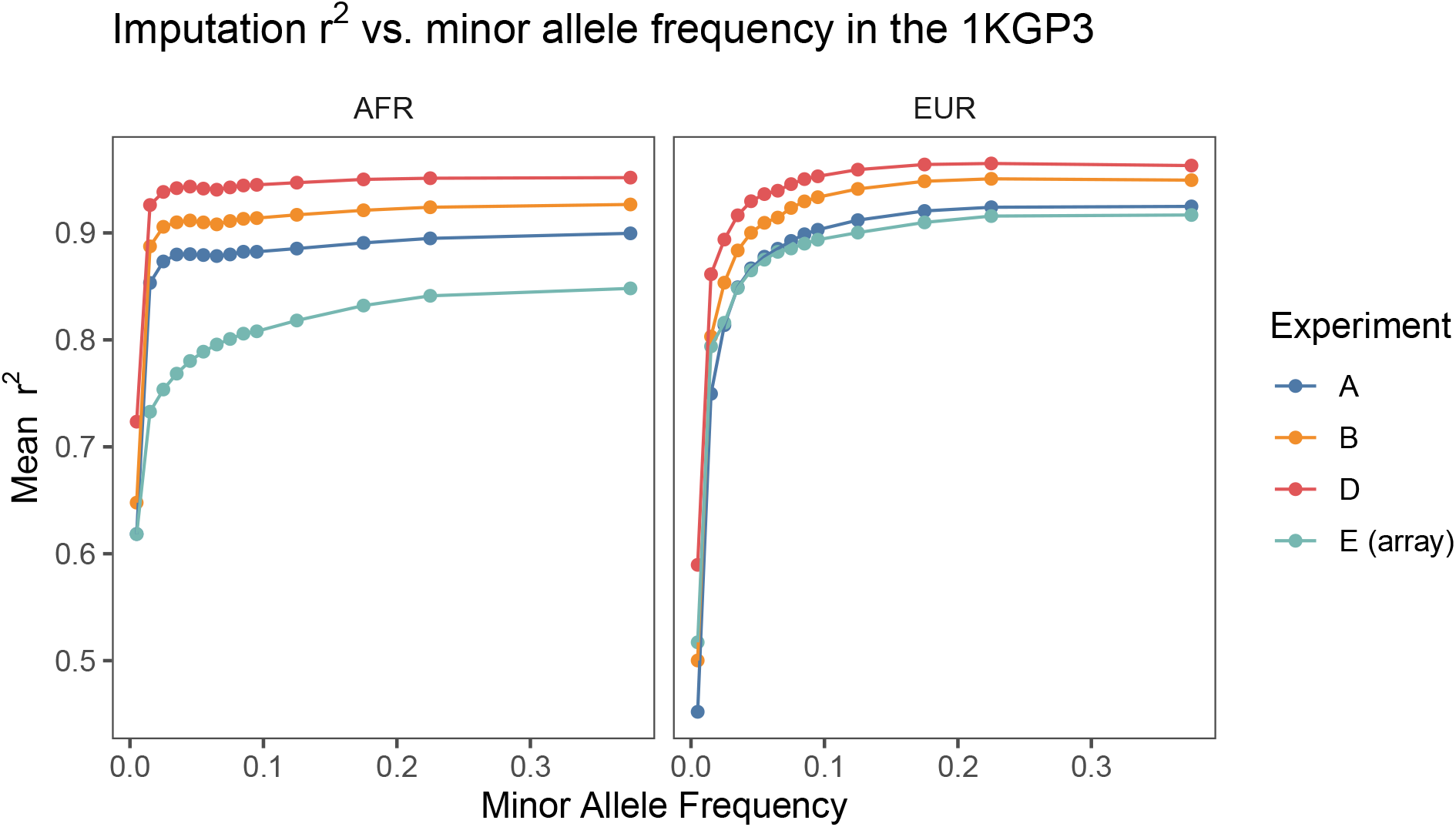
Comparison of imputation quality across experiments for each superpopulation. Variants were binned according to their minor allele frequency in the 1KGP3 and imputation *r*^2^ averaged across variants in each bin. For all experiments the 1KGP3 genotypes were treated as “truth” and imputation *r*^2^ was computed by taking the squared correlation coefficient between the vector of imputed alternate allelic dosages and the truth genotypes. Same results on a log scale are shown in Supplementary Figure 13. Note that imputation performance at low MAF for a given sequencing experiment was often higher in the AFR cohort compared to the EUR cohort.

For the AFR cohorts, we observed that sequence-based experiments uniformly outperformed genotyping arrays for all minor allele frequency bins, while for EUR cohorts the same was true above minor allele frequencies of 0.01 for all but the cohort with the lowest mean effective coverage, experiment A (Supplementary Figure 13).

In particular, at minor allele frequencies of ~ 5% and lower, for a given sequence-based experiment, imputation performance in the AFR cohorts often exceeded performance at similar allele frequencies in the EUR cohorts, a pattern opposite to that which was observed at higher allele frequencies (Supplementary Figure 13, Supplementary Tables 13, 14). This pattern was not observed for the imputed array data, where the imputation *r*^2^s for all allele frequency bins were higher in the EUR cohort than in the AFR cohort.

Similarly to Marchini (2019), we hypothesize that this is due to the fact that at higher allele frequencies, the stronger linkage disequilibrium (LD) structure within European populations dominates and affords more accurate haplotype tagging, whereas at lower allele frequencies, the effects of greater genetic diversity in African populations dominate, resulting in a larger number of possible haplotype combinations and an increased chance that any one rare variant is tagged by at least one of these.

Since low-pass sequencing yields measurements at orders of magnitude more sites than genotyping arrays, it is possible to measure a larger range of these possible haplotypes, which is reflected by this pattern holding only for the sequence-based experiments.

### Polygenic risk scores

Polygenic risk scores were calculated from imputed dosages for each sample in all experiments for breast cancer (BC) and coronary artery disease (CAD) using recent published results in order to evaluate the accuracy of PRS estimation across experiments and assay types (Inouye et al., 2018; Mavaddat et al., 2019). We chose to use weights from these studies because of their demonstrated ability to stratify disease risk beyond conventional risk factors. Of the 1,745,179 variants with nonzero effect sizes in the CAD PRS, 1,738,589 (99.6%) were present in the haplotype reference panel, and 225,667 were directly typed on the GSA. Of the 313 variants with nonzero effect sizes in the BC PRS, all were present in the haplotype reference panel, and 75 were directly typed on the GSA.

We compared the PRS estimates from each sample to the “true” PRS calculated for that cell line from the 1KGP3 genotypes (Supplementary Figure 15). The estimated scores were highly correlated with the true scores for all experiments, and the *r*^2^s for AFR estimates within experiments and across traits were consistently lower than those for their corresponding EUR counterparts (Supplementary Table 16). This difference was particularly pronounced for the CAD PRS, with *r*^2^s ranging from 0.96 − 0.99 across experiments for the EUR samples and *r*^2^s ranging from 0.87 − 0.94 for the AFR samples.

We investigated the relationship between measurement error in PRS (as quantified by the squared error of an estimate) and effective coverage for the sequence-based experiments. Figure 4 shows that the squared error in PRS estimates for all experiments across populations and traits decreases with increasing effective coverage, and that the measurement error in samples sequenced to an effective coverage of ~ 0.5× or higher generally affords lower measurement error than the array-based estimates, with samples sequenced to an effective coverage of more than 1× (experiment D) having an approximately three- to four-fold decrease in squared error for both traits in the AFR cohort and the EUR cohort for CAD, and around the same squared error for the EUR cohort for BC (~ 1.08-fold decrease) (Supplementary Tables 17, 18, 19). Given a superpopulation and trait, the majority of sequence-based PRS estimates had a mean squared error that was significantly (at an *α* = 0.05 level) different than the corresponding array-based PRS estimates (Supp. Table 19).

**Figure 4:**
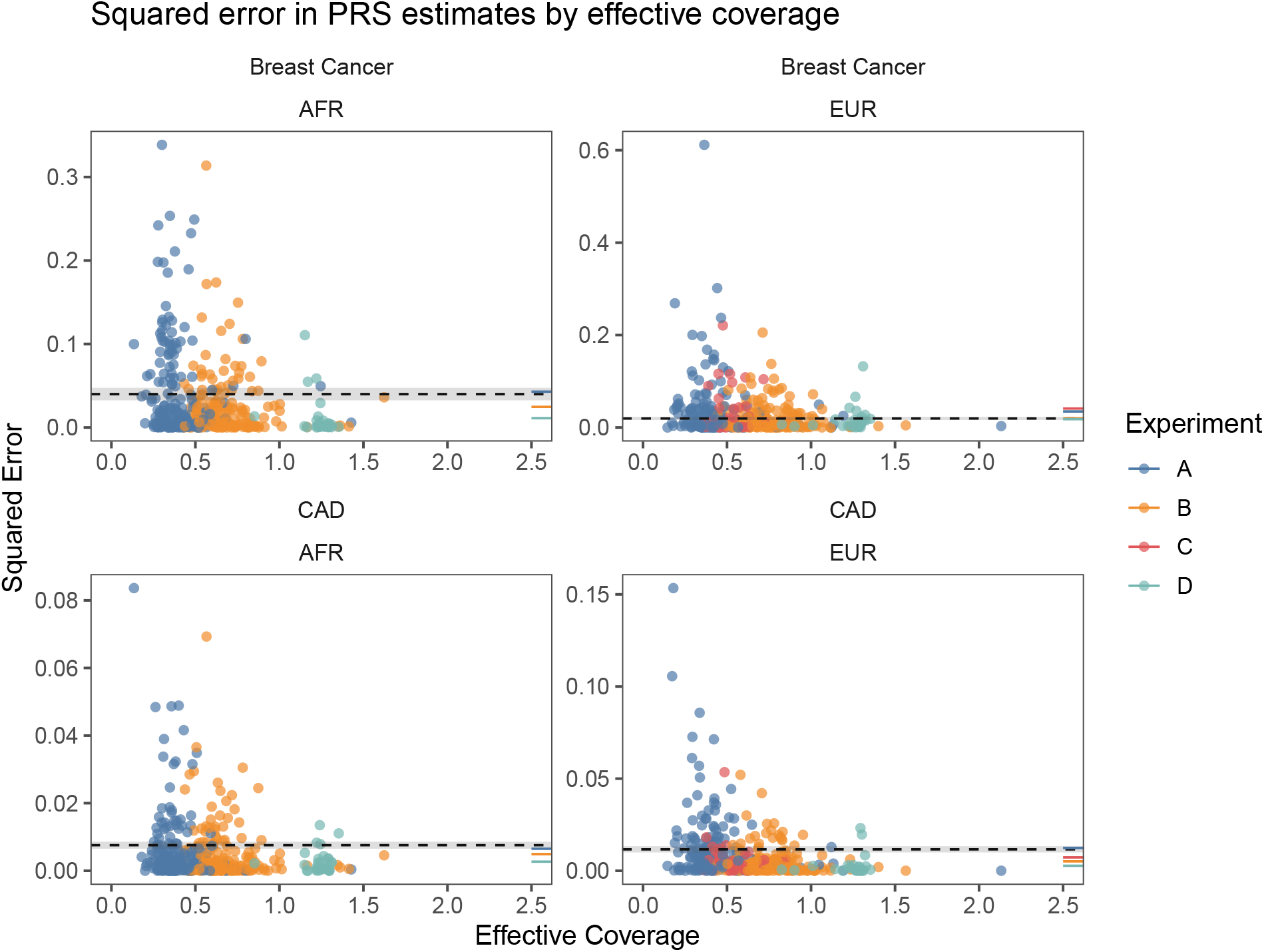
For each imputed sample, we computed a PRS for breast cancer and CAD and calculated the squared error of the estimate compared to the PRS of the “truth” genotypes in the 1KGP3. Each dot represents this squared error for each sequenced sample for a given trait and is colored by which experiment it belongs to. To provide a point of comparison to PRS estimated from imputed array data, we computed the mean squared error for each cell line across array replicates and averaged that across all cell lines. This quantity is represented by the dashed line for each trait and superpopulation along with the standard error of the mean, represented by the shaded regions about each dashed line. The mean squared error for each experiment was calculated in the same way and is rendered as a colored line segment on the rightmost margin of each pane. These results indicate that sequencing at effective coverages of 0.5× or higher generally affords lower measurement error in PRS estimates.

One thing to note is that the measurement error of PRS estimates depends on the quality of genotypes at the variants involved in computing the PRS, which means that for arrays, the measurement error will depend on the proportion of the variants involved that are directly typed versus imputed. Here, a minority of the variants comprising the polygenic risk scores (13% and 24% for CAD and BC respectively) were directly typed on the GSA. Presumably, the measurement error for array-based estimates would be substantially lower in a different situation in which all the variants comprising a PRS are directly typed instead.

## Discussion

The use of dense genotyping arrays followed by imputation to a haplotype reference panel has enabled population-scale genome wide associations studies to become routine in characterizing the genetic architecture of complex traits and disease. An alternative technology is low-pass whole genome sequencing followed by imputation, which has successfully used for a number of problems in statistical and population genetics (Tran et al., 2020; Pasaniuc et al., 2012; Gilly et al., 2019), and which was recently shown to recapitulate comparable disease risk stratification performance using a handful of polygenic risk scores derived from imputed array data from an independent cohort (Homburger et al., 2019).

We introduced the notion of *effective coverage*, a quantity which describes the coverage of a sequenced sample under an ideal sampling process and which is more predictive of imputation accuracy than nominal coverage. We showed that with increasing effective coverage, genotyping concordance and imputation *r*^2^ increase commensurately, while the measurement error of PRS estimates from the imputed genotypes decreases. We note that because the same amount of sequencing coming from different library preparation methods can yield different effective coverages, systematic comparisons of different library preparation methods on this metric is warranted. For instance, the BGIseq library prep used in this experiment contrasts to the Illumina prep in that it is a PCR-free workflow, where every replicated copy generated during DNA nanoball rolling-circle replication comes directly from the native genomic fragment (Patterson et al., 2019; Xia et al., 2019). This has the advantage of reducing propagated amplification error and any potential PCR bias of the template due to selective amplification, well-characterized artifacts of excess PCR.

We observed that at sites of common variation (MAF > 5%), imputation *r*^2^ was consistently and substantially higher in imputed sequence data compared to imputed array data, and in Europeans compared to Africans (Supplementary Table 15). Conversely, at rare variants (MAF < 5%), we observed that imputation *r*^2^ was consistently higher in Africans than Europeans, a result which we hypothesize is due to different aspects of the LD structure in these populations dominating in different regimes of the allele frequency spectrum. A consequence of this observation is that studies of rare variants may (all other things being equal) be more powerful in African-ancestry cohorts as compared to European-ancestry cohorts under some study designs.

We compared PRS estimates for coronary artery disease and breast cancer, and found that the measurement error of PRS estimates decreased with increasing effective coverage for sequenced samples. For CAD, we found that low-pass sequence data yielded consistently lower measurement error in PRS estimates in both the African and European cohorts, with samples sequenced to an effective coverage of ~ 1.2× yielding an approximately three- to four-fold decrease in mean squared error when compared to PRS estimated from the Illumina GSA. The same decrease was observed for BC estimates in the African cohorts, whereas the mean squared error for BC in the European cohort at that effective coverage was around the same as the array estimates (~ 1.08-fold relative decrease).

Since imputation accuracy from genotyping array data is known to depend heavily on the size and composition of the haplotype reference panel used (Schurz et al., 2019), it will be interesting to replicate these results for low-pass sequence across different panels, particularly as extremely large panels (*e.g.*, HRC and TOPmed (McCarthy et al., 2016; Taliun et al., 2019)) as well as panels comprising currently underrepresented populations (*e.g.*, NeuroGAP-Psychosis (Stevenson et al., 2019)) come online.

In the context of understanding of how genetic variation contributes to phenotypic variance, it has become increasingly clear in recent years that a whole-genome approach is necessary due to the fact that the degree of polygenicity underlying the majority of complex traits in humans is quite well-approximated by Fisher’s infinitesimal model (in which variation arises from infinitely many loci of infinitesimal effect), and indeed also that complex traits are mainly driven by non-coding variants (Fisher, 1919; Risch et al., 1999; Barton et al., 2017; Boyle et al., 2017). Consequently, methods to accurately assess genetic variation across the whole genome in diverse populations will be become increasingly essential to ongoing efforts to elucidate the genetic architecture of complex traits, as well as the population genetic processes which gave rise to it (Berg and Coop, 2014; Guo et al., 2018). Our results show that low-pass sequencing is a competitive alternative to genotyping arrays, by affording increased GWAS power and reduced measurement error in the estimation of polygenic risk scores for individuals across multiple populations at a similar price point.

We have discussed the implications of using low-pass sequencing vs. genotyping arrays in the context of PRS estimation and GWAS at length, but there are other, orthogonal considerations that should be taken into account when making a choice between lps and genotyping arrays for practical purposes. For instance, sequence data allows more sensitive detection of copy number and structural variation due to the sheer number of reads that are produced even at extremely low coverages (for intuition, 2.2 million 150bp reads correspond to a coverage of 0.1× for the human genome) (Zhou et al., 2018; Dong et al., 2016). Analysis of mitochondrial count is also straightforward using sequence data, and having sequence data on the mitochondria enables straightforward DNA contamination detection using read count data (Cai et al., 2015b; Krause et al., 2010). Furthermore, the use of sequencing enables analysis of the microbiome and metagenomic profiling by yielding reads deriving from non-human DNA within a given biological sample (Wood et al., 2019).

As low-coverage sequencing becomes increasingly popular as an alternative to genotyping arrays, it is worthwhile to include some discussion on costs and logistical considerations involved in such projects. With sequencing experiments, the two primary factors determining cost are (1) the cost of sequencing and (2) the cost of sample preparation. For instance, the list price of a NovaSeq S4 flow cell as of October 2020 is ≈ 15, 000 US dollars, which corresponds to a sequencing cost of (15,000/1536) = $9.76 per sample for 0.5× sequencing and (15,000/768) = $19.52 per sample for 1× sequencing (NovaSeq Reagent Kits; https://www.illumina.com/products/by-type/sequencing-kits/cluster-gen-sequencing-reagents/novaseq-reagent-kits.html). On the library preparation side, miniaturized/modified workflows currently allow for library preparation costs of under $10/sample (in our specific case, we estimate our costs for the miniaturized Kapa Hyper Prep workflow at around $6/sample). The all-in cost of library preparation and sequencing therefore comes out to be just under $16/sample for 0.5× coverage and just under $26/sample for 1× coverage, comparing favorably with the list price of the Illumina GSA (approximately $49/sample as of November 2020 (Infinium Global Screening Array 24-pack; https://www.illumina.com/products/by-type/microarray-kits/infinium-global-screening.html)).

For future directions, there are methodological advances that could be made in order to fully leverage the information that sequencing provides. Genotyping arrays suffer from ascertainment bias which manifests in two ways: (1) measurements are always made on the same set of loci across the genome, and (2) measurements at a given loci do not allow for novel variant discovery (*i.e.*, you have to know what you are looking for) (Lachance and Tishkoff, 2013; Nielsen, 2004).

Low-pass sequencing by its nature overcomes (1) but whether a problem similar to (2) remains depends on the analytical techniques utilized downstream of actual sequencing. For instance, the current implementation of the imputation tool used here (`loimpute` (Wasik et al., 2019)) does not allow for variants which are novel to the reference panel to be called, thus causing a similar effect in post-imputation sequence data as (2) in unimputed array data. In other words, this means that the effective upper bound in genotyping accuracy is governed by the composition of the reference panel and whether a particular individual’s genetic variation is catalogued therein. A potential methodological improvement would thus be to develop a way to enable novel variant calls at sites with overwhelming read-based evidence.

As research into the genetic architecture of complex disease and traits continues to accelerate, it will become increasingly important for data generation techniques to be as population-agnostic as possible in order to capture global genetic variation in an unbiased manner. Our results demonstrate that low-pass sequencing provides a competitive alternative to genotyping arrays in the context of genome-wide association studies and polygenic risk scoring across diverse populations.

## Methods

### Data generation

Purified genomic DNA (gDNA) from 60 selected individuals of European ancestry and 60 selected individuals of African ancestry from the 1000 Genomes Project Phase 3 was obtained from the NIGMS Human Genetic Cell Repository at the Coriell Institute for Medical Research. Genomic DNA is extracted from immortalized B-lymphocytes and eluted in TE buffer (10mM Tris, pH 8.0/1mM EDTA) for shipping.

To prepare the 120 gDNA samples for sequencing, 30μL from each was plated and diluted to 10ng/μL using 10mM Tris-HCl and sequencing libraries were prepared in triplicate using a miniaturized version of the KAPA HyperPlus kit (Roche, KK8514) with Illumina-compatible KAPA Unique Dual-Indexed Adapters (Roche, KK8727).

Following library preparation, a subset volume of 10μL was pooled from each of the resulting libraries. The pooled libraries were purified with a double-sided size-selection using SPRIselect paramagnetic beads (Beckman Coulter Life Sciences, B23318) to narrow the library fragment size range and remove dimerized adapters from the samples. To characterize the purified pools, the concentration was measured using the Invitrogen Qubit Fluorometer (Thermo Fisher Scientific, Q33238) and the library fragment size was assessed using the Agilent 2100 Bioanalyzer (Agilent, G2939BA) with the High Sensitivity DNA Kit (Agilent, 5067-4626).

Sequencing of purified library pools of 30 and 60 samples was performed to 0.5× coverage and 1× target coverages, respectively, using paired-end 150bp reads on the Illumina HiSeqX platform.

To prepare NA12878 for sequencing 30 times to 1× target coverage, 10μL from the source gDNA sample was aliquoted into 30 separate wells of a 96-well plate and diluted to 10ng/μL using 10mM Tris-HCl. Sequencing libraries were prepared from these diluted samples using a miniaturized version of the KAPA HyperPlus kit (Roche, KK8514) with Illumina-compatible KAPA Unique Dual-Indexed Adapters (Roche, KK8727). The 30 libraries were pooled and purified with a double-sided size-selection using SPRIselect paramagnetic beads (Beckman Coulter Life Sciences, B23318), checked for concentration and fragment size, and sequenced on a single lane of an Illumina HiSeqX flow cell.

A subset of 1μg gDNA from 30 samples selected from each population (60 total) was aliquoted into plates and submitted to BGI Americas Corporation for DNA nanoball library prep and sequencing (DNBseq™) using paired-end 100bp reads on the BGISEQ-500 sequencing instrument.

Sample genotyping was performed using the Infinium Illumina Global Screening Array v3.0 (Illumina, 20030770) by the Broad Institute Genomic Services group. Each of the 120 samples was genotyped in triplicate. To prepare the 360 samples for genotyping, a total mass of 1μg gDNA from each sample was aliquoted into barcode-labeled tubes and submitted to the Broad Institute for processing.

### Quality control

Of the 360 samples in experiment A assayed on the Illumina HiSeqX, 351 passed QC and the remainder failed due to contamination or low read count (less than 0.1× nominal coverage). Of the 360 samples in experiment B assayed on the Illumina HiSeqX, 350 passed QC and the remainder failed due to contamination or low read count (less than 0.1× nominal coverage). Of the 30 samples in experiment B assayed on the Illumina HiSeqX, all 30 passed QC. Of the 60 samples in experiment D sequenced on BGI machines, 58 passed QC and 2 failed due to contamination. Of the 360 samples in experiment E assayed on the Illumina GSA, 358 passed QC and 2 failed due to low call rate (below 97%).

See Supplementary Tables 2, 3 for a more comprehensive breakdown.

### Effective coverage

Consider an idealised sampling process whereby shotgun sequence data is generated for a given sample to a coverage of *λ*. We can then model the number of reads *k* on a site on the genome as a random variable described by a Poisson distribution thus parameterized. Recall that the probability mass function for a Poisson distribution with these parameter is

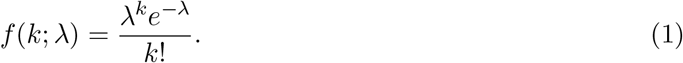

Then the probability that this site is covered by at least one read is

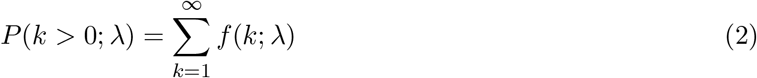

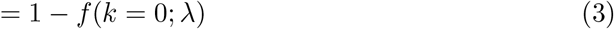

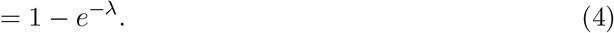

Suppose now that we have a set of *n* such sites on the genome (which in our case represents the *n* sites in a haplotype reference panel) which are all independent and identically distributed. Then the total number of sites *X* covered by at least one read is described by a binomial distribution with *p* = *P* (*k* > 0; *λ*) = 1 − *e^−λ^*, which has an expected value of E[*X*] = *np* = *n*(1 − *e^−λ^*). Defining *f*_covered_ = *X/n* as the *proportion* of sites covered, we have E[*f*_covered_] = (1 − *e^−λ^*).

This quantity describes the expected proportion of sites in a haplotype reference panel covered by at least one read under the assumptions described above, and *f*_covered_ is a quantity which can be empirically computed for any given sample that has been sequenced. The definition of *effective coverage* then follows naturally as that value of *λ* which one obtains when plugging in an observed *f*_covered_ into the relation:

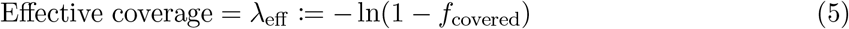

which, as we show in the main text, is more predictive of variant call accuracy than nominal mapped coverage.

### Overall and non-reference concordance

Consider two sets of genotype calls at the same set of *n* sites/variants, with each genotype being coded 0, 1, and 2 for homozygous reference, heterozygous, and homozygous alternate allele. Assume further that there is no missingness in these genotypes. Then at each site, compare the genotype calls between the two callsets — there are nine possible combinations of genotypes ((0, 0), (0, 1), (0, 2), (1, 0), (1, 1), (1, 2), (2, 0), (2, 1), (2, 2)). Each combination can be represented as a cell in a 3×3 table like Table 3, and the total number of each combination can be tallied across all sites. For instance, if the callsets had 100 sites at which both samples were homozygous reference (corresponding to (0, 0)), then at the end of tallying, *a* would be equal to 100 in Table 3.

**Table 3:** Possible combinations of genotype calls between two callsets at a given biallelic site. The diagonal represents concordant calls.

|  | 0 | 1 | 2 |
|---|---|---|---|
| 0 | a | b | c |
| 1 | d | e | f |
| 2 | g | h | i |

Then we define the non-reference concordance (NRC) between the two callsets as

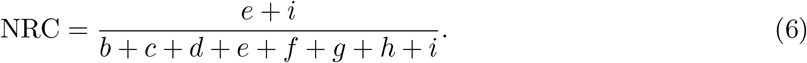

Similarly, the overall concordance is defined as

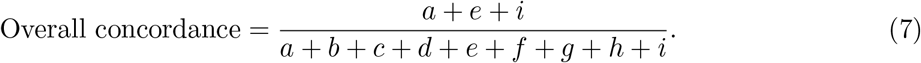

### Selecting samples for representative cohorts

In order to select the representative samples for each experiment for each superpopulation cohort in section, we chose one sample for each cell line for each super population in each experiment so that results compared across assays would concern the exact same samples.

For AFR cell lines, there was at least one replicate in experiments A, B, and E which passed QC for all 60 unique cell lines, so the set of representative samples comprised all 60 cell lines (Supplementary Table 3). For EUR cell lines, there were 57 out of 60 unique cell lines which had at least one replicate in experiments A, B, and E which passed QC, so the set of representative sample comprised a single sample per experiment of these 57 cell lines (Supplementary Table 3).

### Imputation

For the genotyping array data, we used `Eagle v2.4.1` Loh et al. (2016) to perform reference-based phasing for each sample and `minimac4` (Das et al., 2015) for imputation. We received sample-level VCFs with genotypes on the hg19 build of the human reference genome from the Broad Institute; to prepare the samples for phasing and imputation, we filtered out failing probes and duplicate/multiallelic variant probes as marked in the `FILTER` field of the VCFs.

The sequence data were aligned to the `hs37-1kg` reference genome (obtained from NCBI at the following URL: ftp://ftp-trace.ncbi.nih.gov/1000genomes/ftp/technical/reference/human_g1k_v37.fasta.gz) and Gencove’s `loimpute` for imputation. The model underlying Gencove’s `loimpute` has been previously described in the supplementary note to Wasik et al. (2019).

We note that the results of this study should not change in any non-trivial sense if analyses were repeated using build 38 of the human reference genome, as the most substantial differences between the builds are not relevant to imputation of variants on well-characterized regions of the genome (wherein the majority of the variation catalogued by the 1KGP3 callset reside).

To confirm that the qualitative results held when using other imputation software, we repeated the imputation process for the representative cohorts across experiments for Chromosome 21 using `IMPUTE5` for the array data and `GLIMPSE` for the sequence data. The resulting NRC by allele frequency curves showed qualitatively similarly results, with the imputed sequence data showing consistently higher accuracy than the imputed array data across the allele frequency spectrum (Supplementary Figure 12).

All samples were imputed to the 1000 Genomes Phase 3 release haplotype reference panel on hg19 (Consortium et al., 2015) in a leave-one-out manner (such that the cell line the sample belongs to was not in the panel being used). The same was done for the reference-based phasing step for the array data.

Poorly imputed variants were marked as those variants for which the none of the posterior probabilities (the `GP` field in a VCF) for the possible genotypes (hom-ref, heterozygous, hom-alt) was greater than 0.9. In other words, when we refer to “filtered variants”, we are referring to the callset of imputed variants with variants with `max(GP) < 0.9` removed.

## Data access

The raw sequence data generated in this study have been submitted to the NCBI BioProject database (https://www.ncbi.nlm.nih.gov/bioproject/) under accession number PRJNA686136. All raw and processed sequencing and microarray data are also available in a public AWS S3 bucket at s3://gencove-sbir/, accessible also at the following URL: https://gencove-sbir.s3.amazonaws.com/index.html. The code used to perform tertiary analysis, figure generation, and table generation along with the source code for this paper itself as well as the supplementary materials is publicly available at https://gitlab.com/gencove/data-science/presentations-papers-publications/sbir-paper. The `loimpute` software is available at the following URL under a non-commercial license: https://gitlab.com/gencove/loimpute-public.

## Acknowledgements

J.H.L., C.A.M, T.B., and J.K.P. conceived of the study. J.H.L conducted the analyses. C.A.M was responsible for the experimental work and data acquisition. J.H.L., J.K.P., T.B., and C.A.M. contributed to and provided feedback on the manuscript.

Research reported in this publication was supported by a Phase 1 SBIR grant from the NIH (contract number 1R43HG010596-01).

We thank Maria Vazquez for feedback on the study design and operational support of the sequencing and genotyping experiments.

## References

Aird, D., Ross, M. G., Chen, W.-S., Danielsson, M., Fennell, T., Russ, C., Jaffe, D. B., Nusbaum, C., and Gnirke, A., 2011. Analyzing and minimizing PCR amplification bias in Illumina sequencing libraries. Genome biology, 12(2):R18.

Barton, N. H., Etheridge, A. M., and Véber, A., 2017. The infinitesimal model: Definition, derivation, and implications. Theoretical population biology, 118:50–73.

Berg, J. J. and Coop, G., 2014. A population genetic signal of polygenic adaptation. PLoS genetics, 10(8).

Boyle, E. A., Li, Y. I., and Pritchard, J. K., 2017. An expanded view of complex traits: from polygenic to omnigenic. Cell, 169(7):1177–1186.

Cai, N., Bigdeli, T. B., Kretzschmar, W., Li, Y., Liang, J., Song, L., Hu, J., Li, Q., Jin, W., Hu, Z., et al., 2015a. Sparse whole-genome sequencing identifies two loci for major depressive disorder. Nature, 523(7562):588–591.

Cai, N., Li, Y., Chang, S., Liang, J., Lin, C., Zhang, X., Liang, L., Hu, J., Chan, W., Kendler, K. S., et al., 2015b. Genetic control over mtDNA and its relationship to major depressive disorder. Current Biology, 25(24):3170–3177.

Chapman, J. M., Cooper, J. D., Todd, J. A., and Clayton, D. G., 2003. Detecting disease associations due to linkage disequilibrium using haplotype tags: a class of tests and the determinants of statistical power. Human heredity, 56(1-3):18–31.

Consortium,. G. P. et al., 2015. A global reference for human genetic variation. Nature, 526(7571):68–74.

Consortium, W. T. C. C. et al., 2007. Genome-wide association study of 14,000 cases of seven common diseases and 3,000 shared controls. Nature, 447(7145):661.

Das, S., Abecasis, G., and Fuchsberger, C., 2015. Minimac4: A next generation imputation tool for mega reference panels. Abstract 1278W. Presented at the the 65th Annual Meeting of the American Society of Human Genetics, October 7, 2015, Baltimore, MD.

Dong, Z., Zhang, J., Hu, P., Chen, H., Xu, J., Tian, Q., Meng, L., Ye, Y., Wang, J., Zhang, M., et al., 2016. Low-pass whole-genome sequencing in clinical cytogenetics: a validated approach. Genetics in Medicine, 18(9):940–948.

Fisher, R. A., 1919. XV.—The correlation between relatives on the supposition of Mendelian inheritance. Earth and Environmental Science Transactions of the Royal Society of Edinburgh, 52(2):399–433.

Gilly, A., Southam, L., Suveges, D., Kuchenbaecker, K., Moore, R., Melloni, G. E., Hatzikotoulas, K., Farmaki, A.-E., Ritchie, G., Schwartzentruber, J., et al., 2019. Very low-depth whole-genome sequencing in complex trait association studies. Bioinformatics, 35(15):2555–2561.

Guo, J., Wu, Y., Zhu, Z., Zheng, Z., Trzaskowski, M., Zeng, J., Robinson, M. R., Visscher, P. M., and Yang, J., 2018. Global genetic differentiation of complex traits shaped by natural selection in humans. Nature communications, 9(1):1–9.

Harrell, F., 2017. Regression modeling strategies. BIOS, 330:2018.

Homburger, J. R., Neben, C. L., Mishne, G., Zhou, A. Y., Kathiresan, S., and Khera, A. V., 2019. Low coverage whole genome sequencing enables accurate assessment of common variants and calculation of genome-wide polygenic scores. Genome medicine, 11(1):1–12.

Inc., I., 2020a. Infinium Global Screening Array-24 Kit.

Inc., I., 2020b. NovaSeq Reagent Kits.

Inouye, M., Abraham, G., Nelson, C. P., Wood, A. M., Sweeting, M. J., Dudbridge, F., Lai, F. Y., Kaptoge, S., Brozynska, M., Wang, T., et al., 2018. Genomic risk prediction of coronary artery disease in 480,000 adults: implications for primary prevention. Journal of the American College of Cardiology, 72(16):1883–1893.

Jensen, S. E., Charles, J. R., Muleta, K., Bradbury, P. J., Casstevens, T., Deshpande, S. P., Gore, M. A., Gupta, R., Ilut, D. C., Johnson, L., et al., 2020. A sorghum practical haplotype graph facilitates genome-wide imputation and cost-effective genomic prediction. The Plant Genome, 13(1):e20009.

Jones, M. B., Highlander, S. K., Anderson, E. L., Li, W., Dayrit, M., Klitgord, N., Fabani, M. M., Seguritan, V., Green, J., Pride, D. T., et al., 2015. Library preparation methodology can influence genomic and functional predictions in human microbiome research. Proceedings of the National Academy of Sciences, 112(45):14024–14029.

Judson, H. F., 1979. The eighth day of creation. New York, :550.

Krause, J., Briggs, A. W., Kircher, M., Maricic, T., Zwyns, N., Derevianko, A., and Pääbo, S., 2010. A complete mtDNA genome of an early modern human from Kostenki, Russia. Current Biology, 20(3):231–236.

Lachance, J. and Tishkoff, S. A., 2013. SNP ascertainment bias in population genetic analyses: why it is important, and how to correct it. Bioessays, 35(9):780–786.

Li, Y., Willer, C., Sanna, S., and Abecasis, G., 2009. Genotype imputation. Annual review of genomics and human genetics, 10:387–406.

Liu, S., Huang, S., Chen, F., Zhao, L., Yuan, Y., Francis, S. S., Fang, L., Li, Z., Lin, L., Liu, R., et al., 2018. Genomic analyses from non-invasive prenatal testing reveal genetic associations, patterns of viral infections, and Chinese population history. Cell, 175(2):347–359.

Loh, P.-R., Danecek, P., Palamara, P. F., Fuchsberger, C., Reshef, Y. A., Finucane, H. K., Schoenherr, S., Forer, L., McCarthy, S., Abecasis, G. R., et al., 2016. Reference-based phasing using the Haplotype Reference Consortium panel. Nature genetics, 48(11):1443.

Marchini, J., 2019. Haplotype Estimation and Genotype Imputation, chapter 3, pages 87–114. John Wiley & Sons, Ltd.

Marchini, J. and Howie, B., 2010. Genotype imputation for genome-wide association studies. Nature Reviews Genetics, 11(7):499–511.

Mavaddat, N., Michailidou, K., Dennis, J., Lush, M., Fachal, L., Lee, A., Tyrer, J. P., Chen, T.-H., Wang, Q., Bolla, M. K., et al., 2019. Polygenic risk scores for prediction of breast cancer and breast cancer subtypes. The American Journal of Human Genetics, 104(1):21–34.

McCarthy, S., Das, S., Kretzschmar, W., Delaneau, O., Wood, A. R., Teumer, A., Kang, H. M., Fuchsberger, C., Danecek, P., Sharp, K., et al., 2016. A reference panel of 64,976 haplotypes for genotype imputation. Nature genetics, 48(10):1279–1283.

Nielsen, R., 2004. Population genetic analysis of ascertained SNP data. Human genomics, 1(3):218.

Pasaniuc, B., Rohland, N., McLaren, P. J., Garimella, K., Zaitlen, N., Li, H., Gupta, N., Neale, B. M., Daly, M. J., Sklar, P., et al., 2012. Extremely low-coverage sequencing and imputation increases power for genome-wide association studies. Nature genetics, 44(6):631.

Patterson, J., Carpenter, E. J., Zhu, Z., An, D., Liang, X., Geng, C., Drmanac, R., and Wong, G. K.-S., 2019. Impact of sequencing depth and technology on de novo RNA-seq assembly. BMC genomics, 20(1):604.

Pritchard, J. K. and Przeworski, M., 2001. Linkage disequilibrium in humans: models and data. The American Journal of Human Genetics, 69(1):1–14.

Risch, N., Spiker, D., Lotspeich, L., Nouri, N., Hinds, D., Hallmayer, J., Kalaydjieva, L., McCague, P., Dimiceli, S., Pitts, T., et al., 1999. A genomic screen of autism: evidence for a multilocus etiology. The American Journal of Human Genetics, 65(2):493–507.

Rubinacci, S., Ribeiro, D., Hofmeister, R., and Delaneau, O., 2020. Efficient phasing and imputation of low-coverage sequencing data using large reference panels. bioRxiv,.

Schurz, H., Müller, S. J., Van Helden, P. D., Tromp, G., Hoal, E. G., Kinnear, C. J., and Möller, M., 2019. Evaluating the accuracy of imputation methods in a five-way admixed population. Frontiers in genetics, 10:34.

Sella, G. and Barton, N. H., 2019. Thinking about the evolution of complex traits in the era of genome-wide association studies. Annual review of genomics and human genetics, 20:461–493.

Stevenson, A., Akena, D., Stroud, R. E., Atwoli, L., Campbell, M. M., Chibnik, L. B., Kwobah, E., Kariuki, S. M., Martin, A. R., de Menil, V., et al., 2019. Neuropsychiatric Genetics of African Populations-Psychosis (NeuroGAP-Psychosis): a case-control study protocol and GWAS in Ethiopia, Kenya, South Africa and Uganda. BMJ open, 9(2):bmjopen–2018.

Taliun, D., Harris, D. N., Kessler, M. D., Carlson, J., Szpiech, Z. A., Torres, R., Taliun, S. A. G., Corvelo, A., Gogarten, S. M., Kang, H. M., et al., 2019. Sequencing of 53,831 diverse genomes from the NHLBI TOPMed Program. BioRxiv,.

Tran, N. H., Vo, T. B., Tran, N.-T., Trinh, T.-H. N., Pham, H.-A. T., Dao, T. H. T., Nguyen, N. M., Van, Y.-L. T., Tran, V. U., Vu, H. G., et al., 2020. Genetic profiling of Vietnamese population from large-scale genomic analysis of non-invasive prenatal testing data. Scientific reports, 10(1):1–8.

Visscher, P. M., Brown, M. A., McCarthy, M. I., and Yang, J., 2012. Five years of GWAS discovery. The American Journal of Human Genetics, 90(1):7–24.

Walsh, B. and Lynch, M., 2018. Evolution and selection of quantitative traits. Oxford University Press.

Wasik, K., Berisa, T., Pickrell, J. K., Li, J. H., Fraser, D. J., King, K., and Cox, C., 2019. Comparing low-pass sequencing and genotyping for trait mapping in pharmacogenetics. bioRxiv,.

Wetterstrand, K. A., 2019. DNA sequencing costs: Data from the NHGRI Genome Sequencing Program (GSP). National Human Genome Research Institute 2019..

Wood, D. E., Lu, J., and Langmead, B., 2019. Improved metagenomic analysis with Kraken 2. Genome biology, 20(1):257.

Xia, Z., Jiang, Y., Drmanac, R., Shen, H., Liu, P., Li, Z., Chen, F., Jiang, H., Shi, S., Xi, Y., et al., 2019. Advanced Whole Genome Sequencing Using a Complete PCR-free Massively Parallel Sequencing (MPS) Workflow. bioRxiv,.

Yang, J., Benyamin, B., McEvoy, B. P., Gordon, S., Henders, A. K., Nyholt, D. R., Madden, P. A., Heath, A. C., Martin, N. G., Montgomery, G. W., et al., 2010. Common SNPs explain a large proportion of the heritability for human height. Nature genetics, 42(7):565.

Zhou, B., Ho, S. S., Zhang, X., Pattni, R., Haraksingh, R. R., and Urban, A. E., 2018. Whole-genome sequencing analysis of CNV using low-coverage and paired-end strategies is efficient and outperforms array-based CNV analysis. Journal of Medical Genetics, 55(11):735–743.

