## Supplementary Materials for "Low-pass sequencing increases the power of GWAS and decreases measurement error of polygenic risk scores compared to genotyping arrays"

### 1 Supplementary Tables

| Population | Super Population | n |
|---|---|---|
| ASW | AFR | 12 |
| ACB | AFR | 8 |
| ESN | AFR | 8 |
| GWD | AFR | 8 |
| LWK | AFR | 8 |
| MSL | AFR | 8 |
| YRI | AFR | 8 |
| GBR | EUR | 24 |
| IBS | EUR | 12 |
| TSI | EUR | 12 |
| CEU | EUR | 6 |
| FIN | EUR | 6 |

**Supplementary Table 1:** Number of unique cell lines assayed from each population for the EUR and AFR super populations.

| Experiment | Samples | Passing QC | Unique EUR | Unique AFR | Mapped coverage | Eff. Coverage |
|---|---|---|---|---|---|---|
| A | 360 | 351 | 60 | 60 | 0.668±0.318 | 0.412 ± 0.188 |
| B | 360 | 350 | 60 | 60 | 1.248±0.263 | 0.717 ± 0.180 |
| C | 30 | 30 | 1 | 0 | 1.201±0.289 | 0.526 ± 0.112 |
| D | 60 | 58 | 30 | 30 | 1.260±0.0892 | 1.240 ± 0.106 |
| E | 360 | 358 | 60 | 60 | NA | NA |

**Supplementary Table 2:** Breakdown of experiments conducted as described in main text. For experiments A, B, and E, 120 unique cell lines were run in triplicate. For experiment C, NA12878 was run 30x in replicate. For experiment C, a subset of 30 for each superpopulation was run once. Mapped coverage was computed as the number of mapped bases divided by 3.3e9. Coverage is reported as the mean ± standard deviation.

**Supplementary Table 3:** This table shows the number of samples for each cell line in each experiment which passed QC. A dot indicates that the particular cell line was not part of the experiment.

| Cell line | Population | Super population | Exp. A | Exp. B | Exp. C | Exp. D | Exp. E |
|---|---|---|---|---|---|---|---|
| HG01879 | ACB | AFR | 3 | 3 | . | 1 | 3 |
| HG01886 | ACB | AFR | 3 | 3 | . | . | 3 |
| HG01889 | ACB | AFR | 3 | 3 | . | 1 | 3 |
| HG01890 | ACB | AFR | 3 | 3 | . | . | 3 |
| HG02009 | ACB | AFR | 3 | 3 | . | 1 | 3 |
| HG02010 | ACB | AFR | 3 | 3 | . | . | 3 |
| HG02107 | ACB | AFR | 3 | 3 | . | 1 | 3 |
| HG02282 | ACB | AFR | 3 | 3 | . | 1 | 3 |

**Supplementary Table 3:** This table shows the number of samples for each cell line in each experiment which passed QC. A dot indicates that the particular cell line was not part of the experiment. *(continued)*

| Cell line | Population | Super population | Exp. A | Exp. B | Exp. C | Exp. D | Exp. E |
|---|---|---|---|---|---|---|---|
| HG02568 | GWD | AFR | 3 | 3 | . | 1 | 3 |
| HG02759 | GWD | AFR | 3 | 3 | . | 1 | 3 |
| HG02760 | GWD | AFR | 3 | 3 | . | . | 3 |
| HG02878 | GWD | AFR | 3 | 3 | . | 1 | 3 |
| HG02879 | GWD | AFR | 3 | 3 | . | 1 | 3 |
| HG02881 | GWD | AFR | 3 | 3 | . | . | 3 |
| HG02882 | GWD | AFR | 3 | 3 | . | . | 3 |
| HG02922 | ESN | AFR | 3 | 3 | . | 1 | 3 |
| HG02938 | ESN | AFR | 3 | 3 | . | . | 3 |
| HG02941 | ESN | AFR | 3 | 3 | . | 1 | 3 |
| HG02982 | GWD | AFR | 3 | 3 | . | 1 | 3 |
| HG03052 | MSL | AFR | 3 | 3 | . | . | 3 |
| HG03054 | MSL | AFR | 3 | 3 | . | 1 | 3 |
| HG03060 | MSL | AFR | 3 | 3 | . | 1 | 3 |
| HG03063 | MSL | AFR | 3 | 3 | . | 1 | 3 |
| HG03077 | MSL | AFR | 3 | 3 | . | 1 | 3 |
| HG03082 | MSL | AFR | 3 | 3 | . | . | 3 |
| HG03095 | MSL | AFR | 3 | 3 | . | . | 3 |
| HG03127 | ESN | AFR | 3 | 3 | . | 1 | 3 |
| HG03202 | ESN | AFR | 3 | 3 | . | . | 3 |
| HG03298 | ESN | AFR | 3 | 3 | . | 1 | 3 |
| HG03382 | MSL | AFR | 3 | 3 | . | . | 3 |
| HG03520 | ESN | AFR | 3 | 3 | . | 1 | 3 |
| HG03521 | ESN | AFR | 3 | 3 | . | . | 3 |
| NA18504 | YRI | AFR | 3 | 3 | . | . | 3 |
| NA18912 | YRI | AFR | 3 | 3 | . | 1 | 3 |
| NA19017 | LWK | AFR | 3 | 3 | . | 1 | 3 |
| NA19041 | LWK | AFR | 3 | 3 | . | . | 3 |
| NA19119 | YRI | AFR | 3 | 3 | . | . | 3 |
| NA19131 | YRI | AFR | 3 | 3 | . | . | 3 |
| NA19152 | YRI | AFR | 3 | 3 | . | . | 3 |
| NA19204 | YRI | AFR | 3 | 3 | . | . | 3 |
| NA19238 | YRI | AFR | 3 | 3 | . | . | 3 |
| NA19239 | YRI | AFR | 3 | 3 | . | . | 3 |
| NA19307 | LWK | AFR | 3 | 3 | . | . | 3 |
| NA19308 | LWK | AFR | 3 | 3 | . | . | 3 |
| NA19310 | LWK | AFR | 3 | 3 | . | 1 | 3 |

**Supplementary Table 3:** This table shows the number of samples for each cell line in each experiment which passed QC. A dot indicates that the particular cell line was not part of the experiment. *(continued)*

| Cell line | Population | Super population | Exp. A | Exp. B | Exp. C | Exp. D | Exp. E |
|---|---|---|---|---|---|---|---|
| NA19350 | LWK | AFR | 3 | 3 | . | 1 | 3 |
| NA19351 | LWK | AFR | 3 | 3 | . | 1 | 3 |
| NA19471 | LWK | AFR | 3 | 3 | . | 1 | 3 |
| NA19625 | ASW | AFR | 3 | 3 | . | 1 | 3 |
| NA19914 | ASW | AFR | 3 | 3 | . | . | 3 |
| NA19922 | ASW | AFR | 3 | 3 | . | . | 3 |
| NA19923 | ASW | AFR | 3 | 3 | . | 1 | 3 |
| NA19984 | ASW | AFR | 3 | 3 | . | . | 3 |
| NA20298 | ASW | AFR | 3 | 3 | . | 1 | 3 |
| NA20320 | ASW | AFR | 3 | 3 | . | . | 3 |
| NA20351 | ASW | AFR | 3 | 3 | . | . | 3 |
| NA20357 | ASW | AFR | 3 | 3 | . | 1 | 3 |
| NA20359 | ASW | AFR | 3 | 3 | . | . | 3 |
| NA20362 | ASW | AFR | 3 | 3 | . | 1 | 2 |
| NA20412 | ASW | AFR | 3 | 3 | . | . | 3 |
| HG00096 | GBR | EUR | 2 | 0 | . | . | 3 |
| HG00101 | GBR | EUR | 3 | 3 | . | . | 3 |
| HG00102 | GBR | EUR | 3 | 3 | . | 1 | 3 |
| HG00105 | GBR | EUR | 3 | 3 | . | 1 | 3 |
| HG00107 | GBR | EUR | 3 | 3 | . | 1 | 3 |
| HG00108 | GBR | EUR | 3 | 3 | . | . | 3 |
| HG00110 | GBR | EUR | 3 | 3 | . | . | 3 |
| HG00111 | GBR | EUR | 3 | 3 | . | 1 | 3 |
| HG00116 | GBR | EUR | 3 | 3 | . | . | 3 |
| HG00119 | GBR | EUR | 3 | 3 | . | . | 3 |
| HG00131 | GBR | EUR | 3 | 3 | . | 1 | 3 |
| HG00132 | GBR | EUR | 3 | 3 | . | 1 | 3 |
| HG00145 | GBR | EUR | 3 | 3 | . | . | 3 |
| HG00160 | GBR | EUR | 3 | 3 | . | 1 | 3 |
| HG00234 | GBR | EUR | 3 | 3 | . | . | 3 |
| HG00239 | GBR | EUR | 3 | 3 | . | 1 | 3 |
| HG00240 | GBR | EUR | 3 | 3 | . | . | 3 |
| HG00242 | GBR | EUR | 3 | 3 | . | 1 | 3 |
| HG00244 | GBR | EUR | 3 | 3 | . | . | 3 |
| HG00250 | GBR | EUR | 3 | 3 | . | 1 | 3 |
| HG00251 | GBR | EUR | 3 | 3 | . | 1 | 3 |
| HG00255 | GBR | EUR | 3 | 3 | . | . | 3 |

**Supplementary Table 3:** This table shows the number of samples for each cell line in each experiment which passed QC. A dot indicates that the particular cell line was not part of the experiment. *(continued)*

| Cell line | Population | Super population | Exp. A | Exp. B | Exp. C | Exp. D | Exp. E |
|---|---|---|---|---|---|---|---|
| HG00262 | GBR | EUR | 3 | 3 | . | . | 3 |
| HG00268 | FIN | EUR | 3 | 3 | . | 1 | 3 |
| HG00330 | FIN | EUR | 3 | 3 | . | . | 3 |
| HG00360 | FIN | EUR | 3 | 3 | . | . | 3 |
| HG00364 | FIN | EUR | 3 | 3 | . | 1 | 3 |
| HG00365 | FIN | EUR | 2 | 3 | . | 0 | 3 |
| HG00384 | FIN | EUR | 3 | 3 | . | . | 3 |
| HG01500 | IBS | EUR | 3 | 3 | . | 1 | 3 |
| HG01697 | IBS | EUR | 3 | 3 | . | . | 3 |
| HG01699 | IBS | EUR | 3 | 3 | . | 1 | 3 |
| HG01756 | IBS | EUR | 3 | 3 | . | 1 | 3 |
| HG01757 | IBS | EUR | 3 | 3 | . | . | 3 |
| HG01761 | IBS | EUR | 3 | 3 | . | . | 3 |
| HG01762 | IBS | EUR | 3 | 3 | . | 1 | 3 |
| HG01770 | IBS | EUR | 3 | 3 | . | 1 | 3 |
| HG01771 | IBS | EUR | 3 | 3 | . | . | 3 |
| HG01777 | IBS | EUR | 2 | 3 | . | 1 | 3 |
| HG01781 | IBS | EUR | 3 | 3 | . | 1 | 3 |
| HG01784 | IBS | EUR | 3 | 3 | . | . | 3 |
| HG02215 | GBR | EUR | 3 | 3 | . | 1 | 3 |
| NA06985 | CEU | EUR | 3 | 3 | . | . | 3 |
| NA11930 | CEU | EUR | 3 | 3 | . | . | 3 |
| NA12342 | CEU | EUR | 3 | 3 | . | 1 | 3 |
| NA12842 | CEU | EUR | 3 | 3 | . | 1 | 3 |
| NA12843 | CEU | EUR | 3 | 3 | . | . | 2 |
| NA12878 | CEU | EUR | 3 | 3 | 30 | 1 | 3 |
| NA20502 | TSI | EUR | 3 | 3 | . | 1 | 3 |
| NA20503 | TSI | EUR | 3 | 3 | . | . | 3 |
| NA20507 | TSI | EUR | 3 | 3 | . | 1 | 3 |
| NA20513 | TSI | EUR | 3 | 2 | . | 1 | 3 |
| NA20514 | TSI | EUR | 3 | 3 | . | . | 3 |
| NA20798 | TSI | EUR | 3 | 3 | . | . | 3 |
| NA20802 | TSI | EUR | 0 | 0 | . | 0 | 3 |
| NA20803 | TSI | EUR | 3 | 3 | . | . | 3 |
| NA20804 | TSI | EUR | 3 | 3 | . | . | 3 |
| NA20805 | TSI | EUR | 3 | 3 | . | 1 | 3 |
| NA20821 | TSI | EUR | 3 | 3 | . | . | 3 |
| NA20822 | TSI | EUR | 0 | 0 | . | . | 3 |

| Coverage | Super Pop. | Experiment A | B | D |
|---|---|---|---|---|
| Nominal | AFR | $0.6409150 \pm 0.11745997$ | $1.2459427 \pm 0.24100086$ | $1.2564230 \pm 0.08498687$ |
| Nominal | EUR | $0.7280792 \pm 0.51483707$ | $1.2149407 \pm 0.17149908$ | $1.2641167 \pm 0.09484703$ |
| Effective | AFR | $0.3711961 \pm 0.08585635$ | $0.6690530 \pm 0.16714982$ | $1.2362189 \pm 0.08771836$ |
| Effective | EUR | $0.4739798 \pm 0.29770421$ | $0.7507368 \pm 0.15740140$ | $1.2345972 \pm 0.12378594$ |

**Supplementary Table 4:** Nominal and effective coverages (each cell is the mean ± standard deviation) of the representative cohorts for each superpopulation for each experiment.

| Super Population | A | B | D | E |
|---|---|---|---|---|
| AFR | 0.9455802 | 0.9533416 | 0.9609383 | 0.9213722 |
| EUR | 0.9625844 | 0.9694407 | 0.9730607 | 0.9663448 |

**Supplementary Table 5:** Mean NRC for filtered SNPs by experiment and super population.

| Super Population | A | B | D | E |
|---|---|---|---|---|
| AFR | 0.8577369 | 0.8737019 | 0.8875245 | 0.7998131 |
| EUR | 0.8661640 | 0.8823104 | 0.8907714 | 0.8548517 |

**Supplementary Table 6:** Mean NRC for unfiltered indels by experiment and super population.

| Super Population | A | B | D | E |
|---|---|---|---|---|
| AFR | 0.9255645 | 0.9286473 | 0.9303212 | 0.9144183 |
| EUR | 0.9379047 | 0.9397931 | 0.9389366 | 0.9480481 |

**Supplementary Table 7:** Mean NRC for filtered indels by experiment and super population.

| Super Population | A | B | D | E |
|---|---|---|---|---|
| AFR | 0.9943481 | 0.9955892 | 0.9967395 | 0.9902573 |
| EUR | 0.9964275 | 0.9974171 | 0.9979954 | 0.9956762 |

**Supplementary Table 8:** Mean overall concordance for unfiltered SNPs by experiment and super population.

| Super Population | A | B | D | E |
|---|---|---|---|---|
| AFR | 0.9972752 | 0.9975687 | 0.9978953 | 0.9966893 |
| EUR | 0.9984646 | 0.9986824 | 0.9988119 | 0.9987583 |

**Supplementary Table 9:** Mean overall concordance for filtered SNPs by experiment and super population.

| Super Population | A | B | D | E |
|---|---|---|---|---|
| AFR | 0.9715808 | 0.9748666 | 0.9776772 | 0.9595724 |
| EUR | 0.9765626 | 0.9794674 | 0.9809911 | 0.9746278 |

**Supplementary Table 10:** Mean overall concordance for unfiltered indels by experiment and super population.

| Super Population | A | B | D | E |
|---|---|---|---|---|
| AFR | 0.9873308 | 0.9873972 | 0.9872877 | 0.9878819 |
| EUR | 0.9909231 | 0.9907706 | 0.9903947 | 0.9933415 |

**Supplementary Table 11:** Mean overall concordance for filtered indels by experiment and super population.

| Super Population | Non-Reference Allele Frequency | A | B | D | E |
|---|---|---|---|---|---|
| AFR | 0.005 | 0.6330749 | 0.6624347 | 0.6915782 | 0.5141283 |
| AFR | 0.015 | 0.8442763 | 0.8780813 | 0.9090730 | 0.6733742 |
| AFR | 0.025 | 0.8700104 | 0.9022795 | 0.9304975 | 0.7055604 |
| AFR | 0.035 | 0.8821727 | 0.9126826 | 0.9396327 | 0.7272472 |
| AFR | 0.045 | 0.8860119 | 0.9161137 | 0.9420034 | 0.7417269 |
| AFR | 0.055 | 0.8886665 | 0.9181082 | 0.9436521 | 0.7598017 |
| AFR | 0.065 | 0.8887966 | 0.9182220 | 0.9431593 | 0.7695656 |
| AFR | 0.075 | 0.8909316 | 0.9203547 | 0.9451238 | 0.7825318 |
| AFR | 0.085 | 0.8936551 | 0.9226183 | 0.9467729 | 0.7930463 |
| AFR | 0.095 | 0.8946790 | 0.9232170 | 0.9478058 | 0.8001786 |
| AFR | 0.125 | 0.8971092 | 0.9253967 | 0.9497405 | 0.8178357 |
| AFR | 0.175 | 0.9011986 | 0.9288376 | 0.9528010 | 0.8372098 |
| AFR | 0.225 | 0.9053344 | 0.9316063 | 0.9543433 | 0.8457356 |
| AFR | 0.375 | 0.9152775 | 0.9377851 | 0.9583135 | 0.8622945 |
| AFR | 0.750 | 0.9537986 | 0.9651469 | 0.9778491 | 0.9234951 |
| EUR | 0.005 | 0.4086514 | 0.4603088 | 0.5030438 | 0.3781868 |
| EUR | 0.015 | 0.7123895 | 0.7756373 | 0.8225086 | 0.6822138 |
| EUR | 0.025 | 0.7869669 | 0.8376289 | 0.8718883 | 0.7090553 |
| EUR | 0.035 | 0.8324885 | 0.8762228 | 0.9022587 | 0.7583118 |
| EUR | 0.045 | 0.8568614 | 0.8966524 | 0.9190304 | 0.7934836 |
| EUR | 0.055 | 0.8730475 | 0.9100724 | 0.9302962 | 0.8159317 |
| EUR | 0.065 | 0.8803156 | 0.9153765 | 0.9352682 | 0.8298032 |
| EUR | 0.075 | 0.8877140 | 0.9226817 | 0.9416631 | 0.8423751 |
| EUR | 0.085 | 0.8968696 | 0.9296256 | 0.9480110 | 0.8555732 |
| EUR | 0.095 | 0.8986139 | 0.9314749 | 0.9493600 | 0.8575277 |
| EUR | 0.125 | 0.9070911 | 0.9388980 | 0.9558629 | 0.8739273 |
| EUR | 0.175 | 0.9157642 | 0.9453105 | 0.9612076 | 0.8899715 |
| EUR | 0.225 | 0.9207281 | 0.9484181 | 0.9628728 | 0.9026118 |
| EUR | 0.375 | 0.9302607 | 0.9533038 | 0.9657107 | 0.9143037 |
| EUR | 0.750 | 0.9650752 | 0.9762219 | 0.9829335 | 0.9589904 |

**Supplementary Table 12:** Mean non-reference concordance by minor allele frequency bin in 1KGP3 for EUR and AFR cohorts by experiment. The values in the Minor Allele Frequency column denote the midpoint of the respective bin. These are the values plotted in Figure 2

| Minor Allele Frequency | A | D | B | E |
|---|---|---|---|---|
| 0.005 | 0.6184364 | 0.7234881 | 0.6477770 | 0.6183798 |
| 0.015 | 0.8532266 | 0.9261001 | 0.8874289 | 0.7327668 |
| 0.025 | 0.8733104 | 0.9383263 | 0.9057120 | 0.7535728 |
| 0.035 | 0.8797922 | 0.9418486 | 0.9099334 | 0.7684120 |
| 0.045 | 0.8801383 | 0.9433202 | 0.9116183 | 0.7802390 |
| 0.055 | 0.8792989 | 0.9414425 | 0.9098476 | 0.7889383 |
| 0.065 | 0.8783703 | 0.9403997 | 0.9080035 | 0.7956326 |
| 0.075 | 0.8798552 | 0.9425548 | 0.9111161 | 0.8008224 |
| 0.085 | 0.8824059 | 0.9443458 | 0.9131975 | 0.8058023 |
| 0.095 | 0.8823828 | 0.9449659 | 0.9138677 | 0.8079947 |
| 0.125 | 0.8853671 | 0.9469828 | 0.9169203 | 0.8181209 |
| 0.175 | 0.8906883 | 0.9500326 | 0.9211740 | 0.8320627 |
| 0.225 | 0.8948852 | 0.9511481 | 0.9239985 | 0.8411895 |
| 0.375 | 0.8996063 | 0.9517120 | 0.9265984 | 0.8481170 |

**Supplementary Table 13:** Mean $r^2$s by minor allele frequency bin in 1KGP3 for the AFR cohort by experiment. The values in the Minor Allele Frequency column denote the midpoint of the respective bin. The bin edges are therefore at 0, 0.01, 0.02, 0.03, 0.04, 0.05, 0.06, 0.07, 0.08, 0.09, 0.1, 0.15, 0.2, 0.25, 0.5. These are the values plotted in the left pane of Figure 3.

| Minor Allele Frequency | A | D | B | E |
|---|---|---|---|---|
| 0.005 | 0.4521868 | 0.5895981 | 0.5001920 | 0.5171781 |
| 0.015 | 0.7495764 | 0.8613694 | 0.8030856 | 0.7939501 |
| 0.025 | 0.8137453 | 0.8938028 | 0.8534747 | 0.8159978 |
| 0.035 | 0.8491713 | 0.9164598 | 0.8836018 | 0.8484690 |
| 0.045 | 0.8668877 | 0.9296342 | 0.9001195 | 0.8649535 |
| 0.055 | 0.8777631 | 0.9362962 | 0.9094547 | 0.8749669 |
| 0.065 | 0.8851983 | 0.9394706 | 0.9143465 | 0.8822995 |
| 0.075 | 0.8925096 | 0.9456321 | 0.9233629 | 0.8854623 |
| 0.085 | 0.8987583 | 0.9503176 | 0.9294991 | 0.8899616 |
| 0.095 | 0.9029525 | 0.9529296 | 0.9333947 | 0.8936177 |
| 0.125 | 0.9119380 | 0.9591721 | 0.9411317 | 0.9002847 |
| 0.175 | 0.9205362 | 0.9639785 | 0.9482848 | 0.9098169 |
| 0.225 | 0.9240213 | 0.9649426 | 0.9505899 | 0.9157568 |
| 0.375 | 0.9247924 | 0.9629698 | 0.9493630 | 0.9167748 |

**Supplementary Table 14:** Mean $r^2$s by minor allele frequency in 1KGP3 for the EUR cohort by experiment. The values in the Minor Allele Frequency column denote the midpoint of the respective bin. The bin edges are therefore at 0, 0.01, 0.02, 0.03, 0.04, 0.05, 0.06, 0.07, 0.08, 0.09, 0.1, 0.15, 0.2, 0.25, 0.5. These are the values plotted in the right pane of Figure 3.

| Super population | Experiment A | B | D | E |
|---|---|---|---|---|
| AFR | 0.8916914 | 0.9208044 | 0.9488131 | 0.8301759 |
| EUR | 0.9159033 | 0.9429037 | 0.9595872 | 0.9073543 |

**Supplementary Table 15:** Mean imputation $r^2$s for each representative cohort for each experiment by superpopulation cohort at biallelic SNPs above 5% minor allele frequency in the 1KGP3.

| Super population | Trait | Experiment A | B | D | E |
|---|---|---|---|---|---|
| AFR | Breast Cancer | 0.8867460 | 0.9353159 | 0.9745389 | 0.8943234 |
| EUR | Breast Cancer | 0.9053879 | 0.9470577 | 0.9697582 | 0.9486035 |
| AFR | CAD | 0.8924037 | 0.9195413 | 0.9379679 | 0.8725896 |
| EUR | CAD | 0.9569958 | 0.9832295 | 0.9894414 | 0.9720993 |

**Supplementary Table 16:** Squared Pearson correlation coefficient ($r^2$) between PRS estimates from imputed dosages vs. true PRS as calculated off 1KGP3 genotypes by superpopulation and trait.

| Super population | Trait | MSE | SE of MSE |
|---|---|---|---|
| AFR | Breast Cancer | 0.0399702 | 0.0074939 |
| EUR | Breast Cancer | 0.0195021 | 0.0034662 |
| AFR | CAD | 0.0075706 | 0.0010238 |
| EUR | CAD | 0.0116090 | 0.0016780 |

**Supplementary Table 17:** MSE in array-based PRS estimates and the standard error of the MSE. The MSE is calculated by first taking the average of the squared error in PRS estimates among replicates of a given cell line and trait, and then averaging over all cell lines within a population for a given trait.

| Experiment | Trait | Super population | MSE | SE of MSE | MSE fold decrease |
|---|---|---|---|---|---|
| A | Breast Cancer | AFR | 0.0429071 | 0.0054744 | 0.9315517 |
| A | Breast Cancer | EUR | 0.0346656 | 0.0058626 | 0.5625766 |
| B | Breast Cancer | AFR | 0.0247107 | 0.0031849 | 1.6175279 |
| B | Breast Cancer | EUR | 0.0197314 | 0.0026004 | 0.9883759 |
| C | Breast Cancer | EUR | 0.0410636 | NA | 0.4749238 |
| D | Breast Cancer | AFR | 0.0112054 | 0.0043408 | 3.5670630 |
| D | Breast Cancer | EUR | 0.0179538 | 0.0050838 | 1.0862357 |
| A | CAD | AFR | 0.0065103 | 0.0010025 | 1.1628575 |
| A | CAD | EUR | 0.0124154 | 0.0016935 | 0.9350478 |
| B | CAD | AFR | 0.0049382 | 0.0007701 | 1.5330638 |
| B | CAD | EUR | 0.0051197 | 0.0006327 | 2.2674925 |
| C | CAD | EUR | 0.0072507 | NA | 1.6010815 |
| D | CAD | AFR | 0.0027052 | 0.0006232 | 2.7985326 |
| D | CAD | EUR | 0.0027173 | 0.0010522 | 4.2722023 |

**Supplementary Table 18:** Mean squared error (MSE) in sequence-based PRS estimates and the standard error (SE) of the MSE. The MSE is calculated by first taking the average of the squared error in PRS estimates among replicates of a given cell line and trait, and then averaging over all cell lines within a population for a given trait. MSE fold decrease is the $x$-fold decrease in MSE compared to the array counterpart. For instance, the MSE for experiment D's CAD PRS estimates for the AFR cohort was approximately four-fold the MSE of the array-based AFR CAD estimates (Supplementary Table 17).

#### 2 Figures

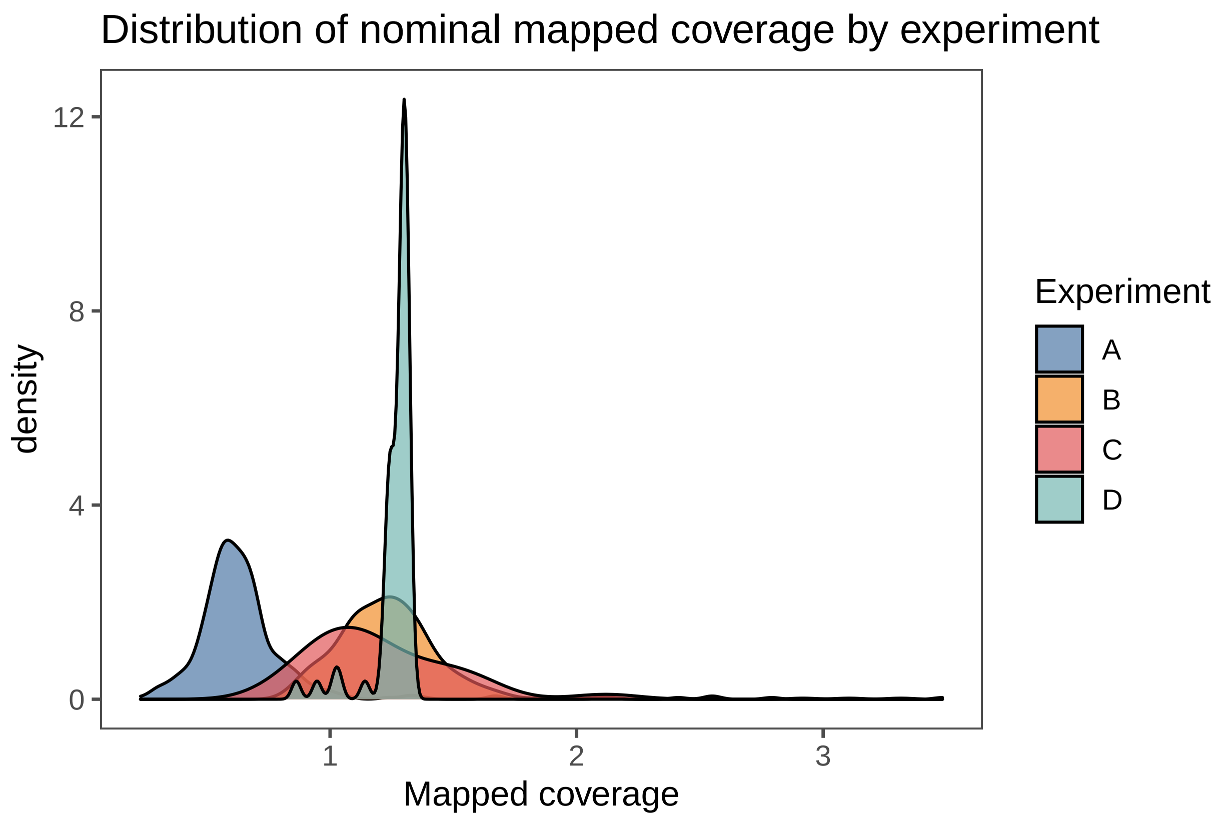


**Supplementary Figure 1:** Distribution of mapped coverages by experiment.

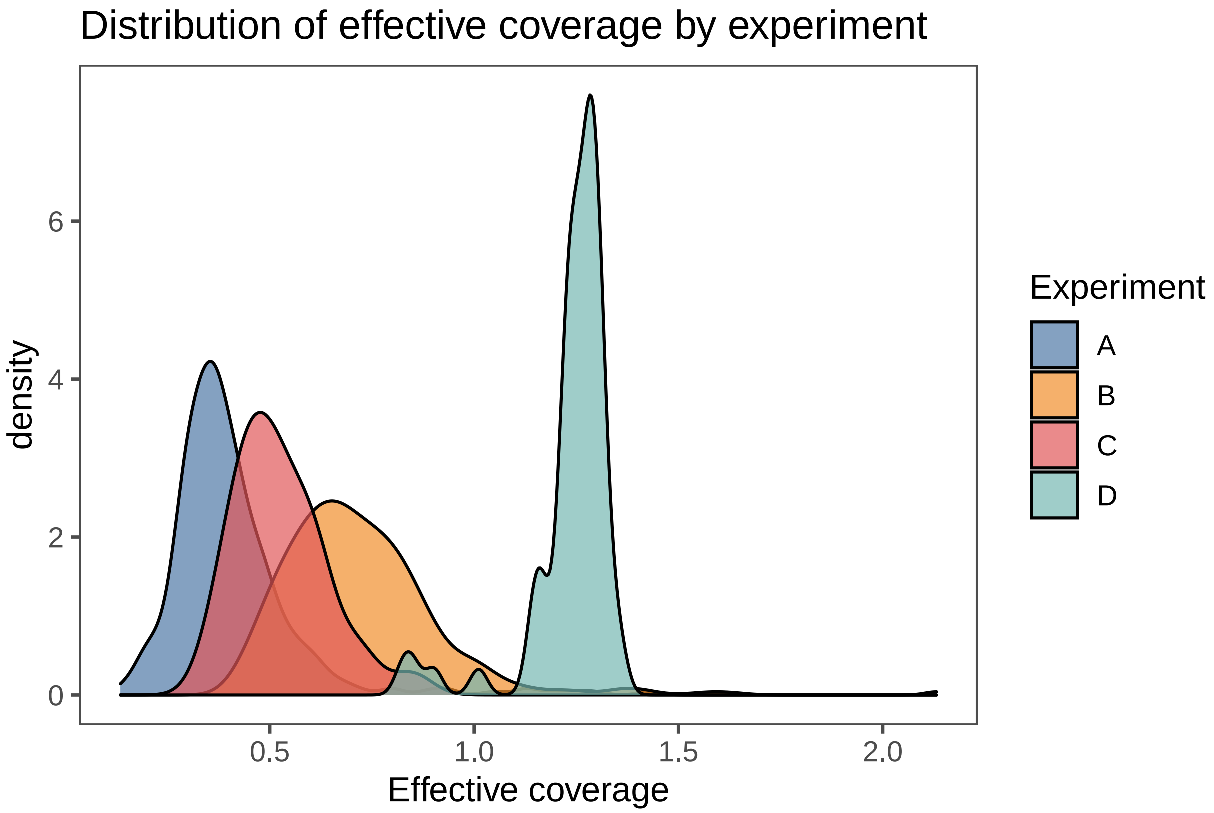


**Supplementary Figure 2:** Distribution of effective coverages by experiment.

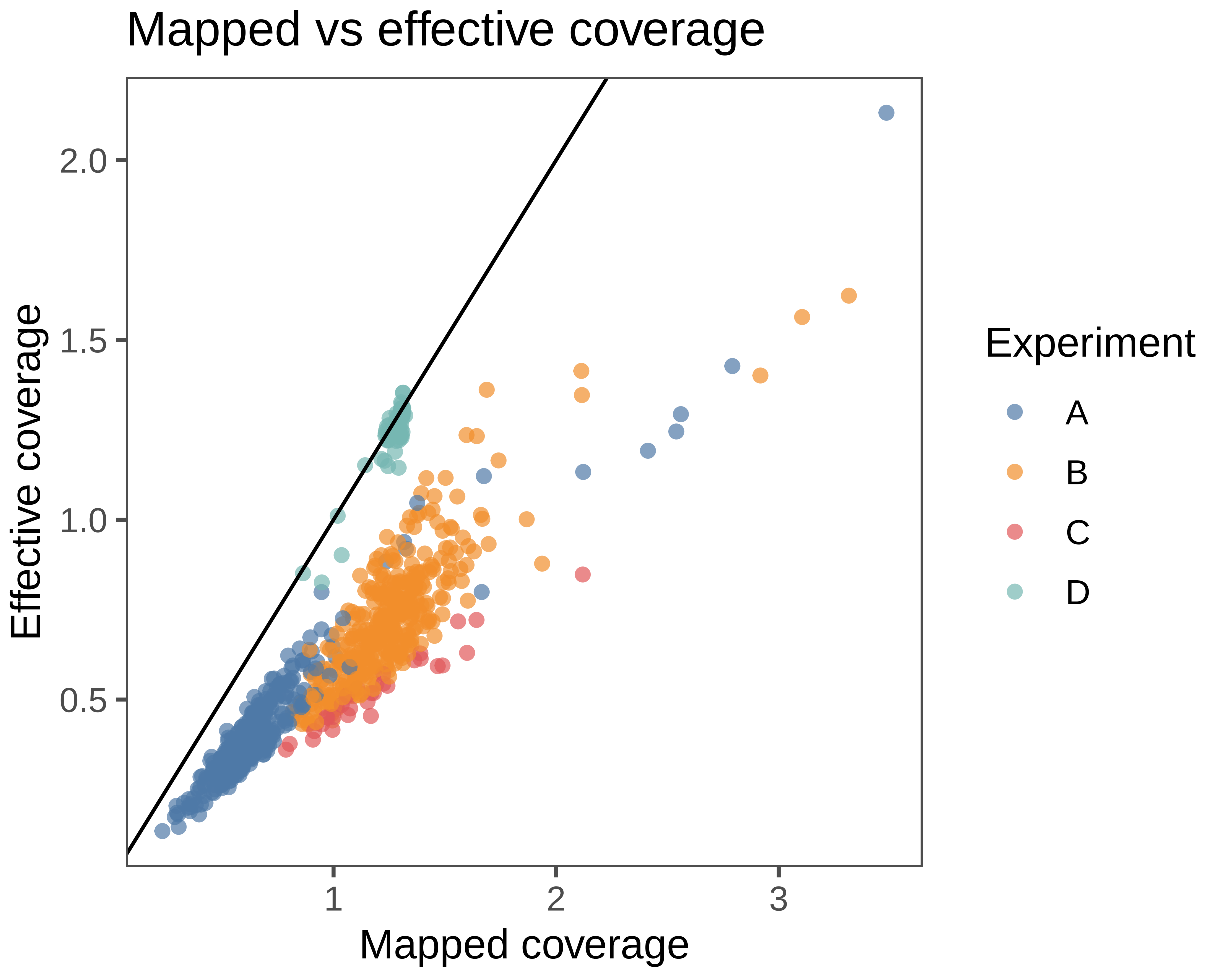


**Supplementary Figure 3:** Mapped deduplicated coverage vs. effective coverage as described in main manuscript.

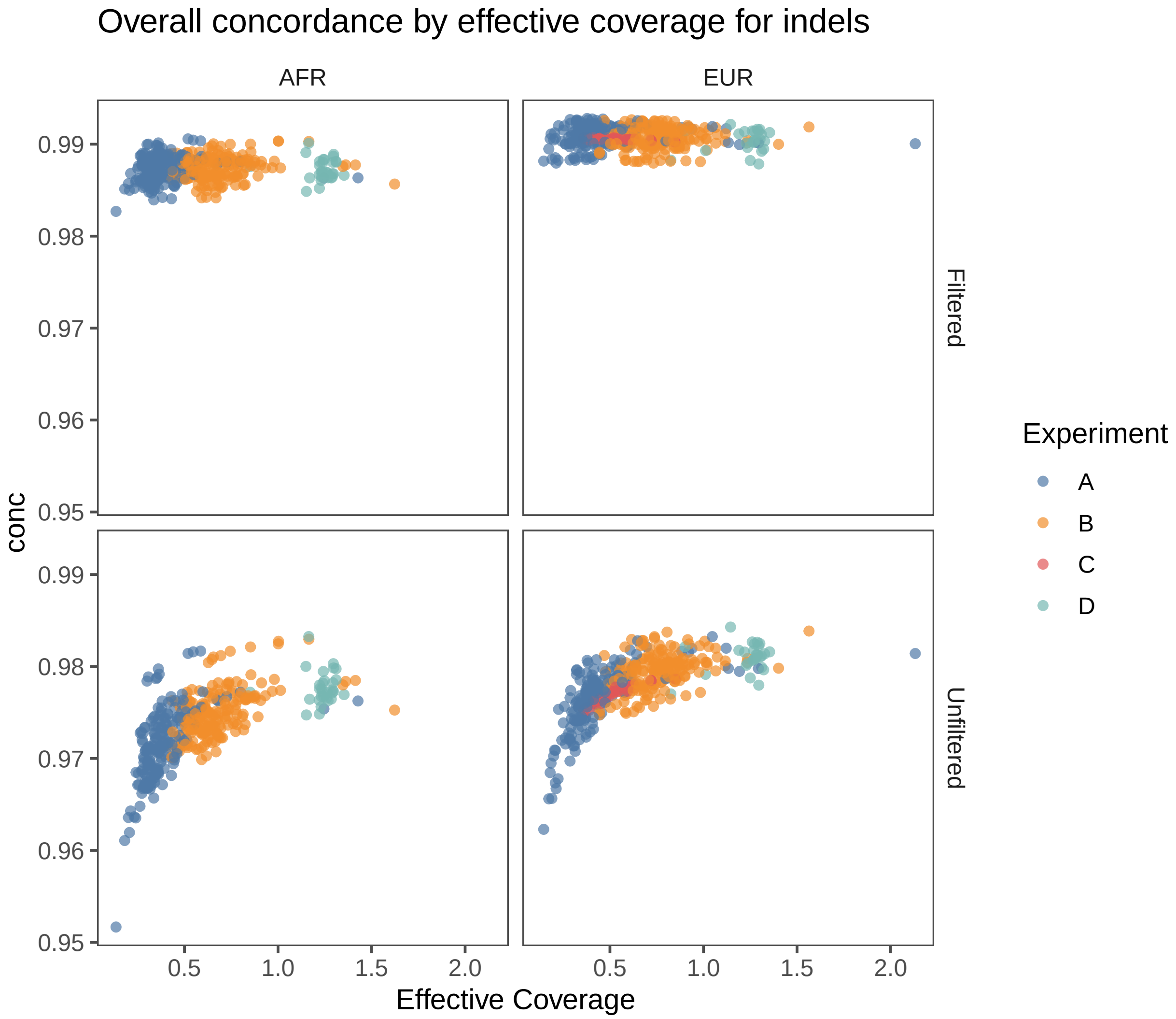


**Supplementary Figure 4:** Overall concordance for SNPs by effective coverage for both filtered and unfiltered variant calls.

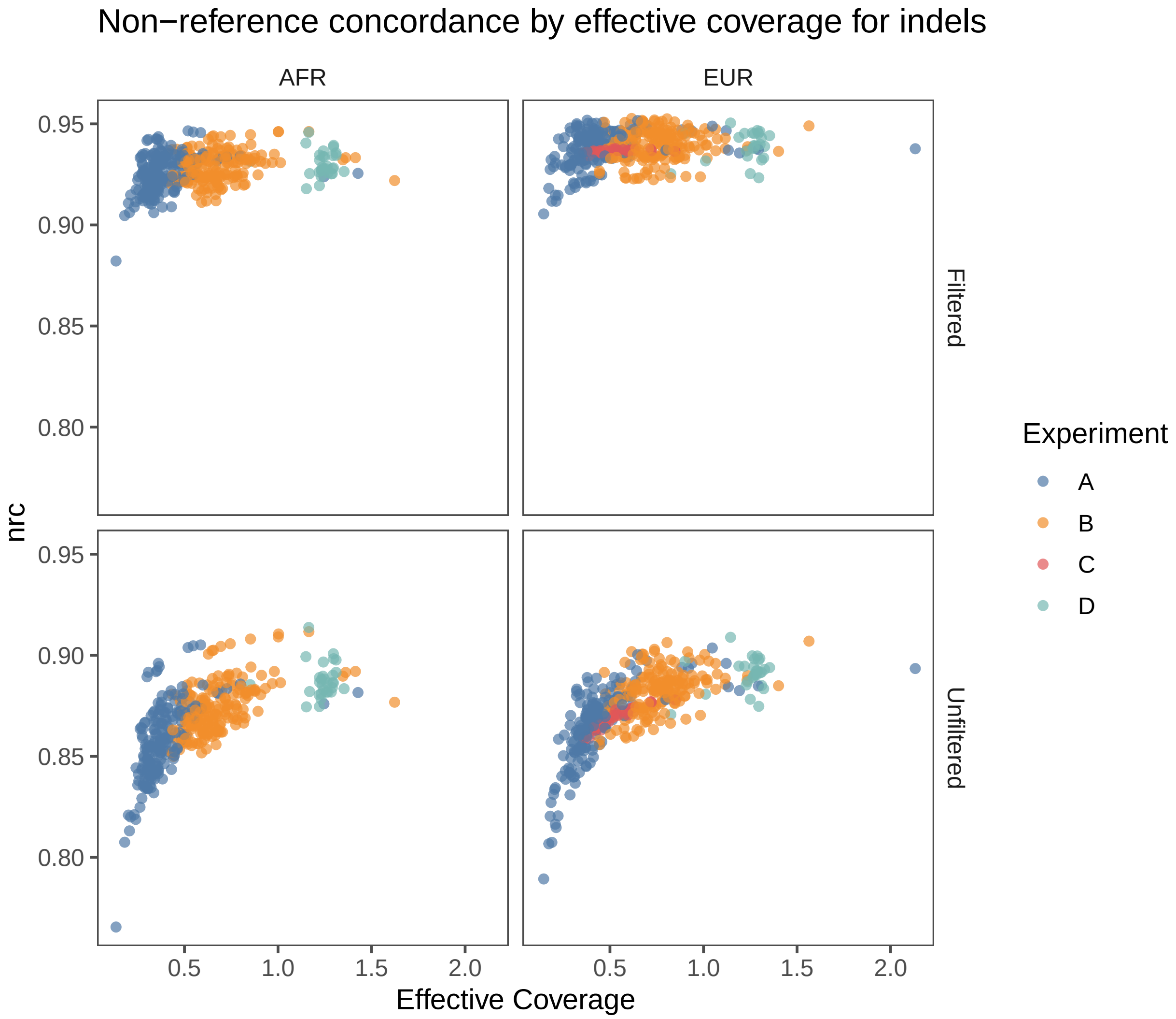


**Supplementary Figure 5:** Non-reference concordance for indels by effective coverage for both filtered and unfiltered variant calls.

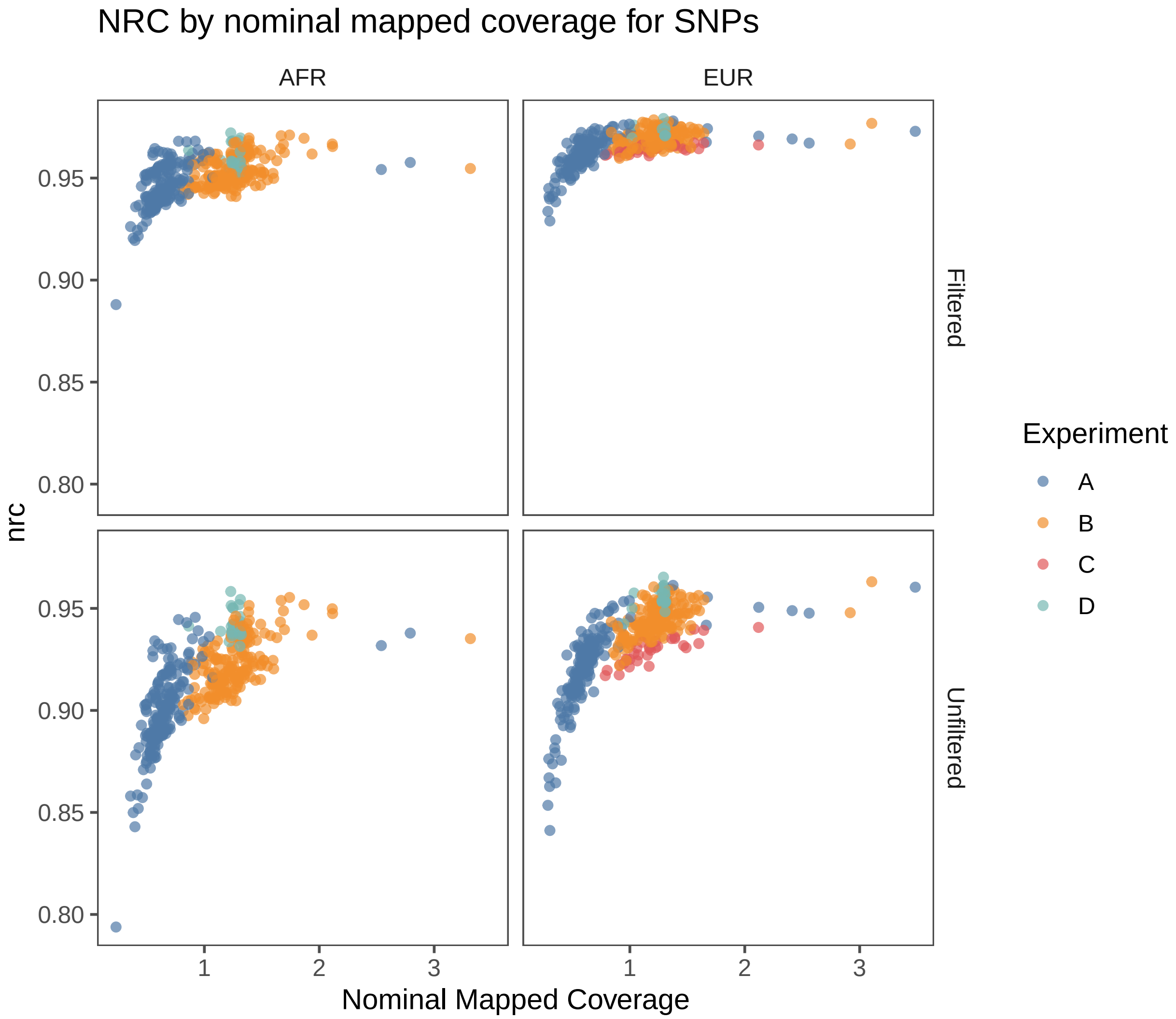


**Supplementary Figure 6:** Non-reference concordance for SNPs by nominal coverage for both filtered and unfiltered variant calls.

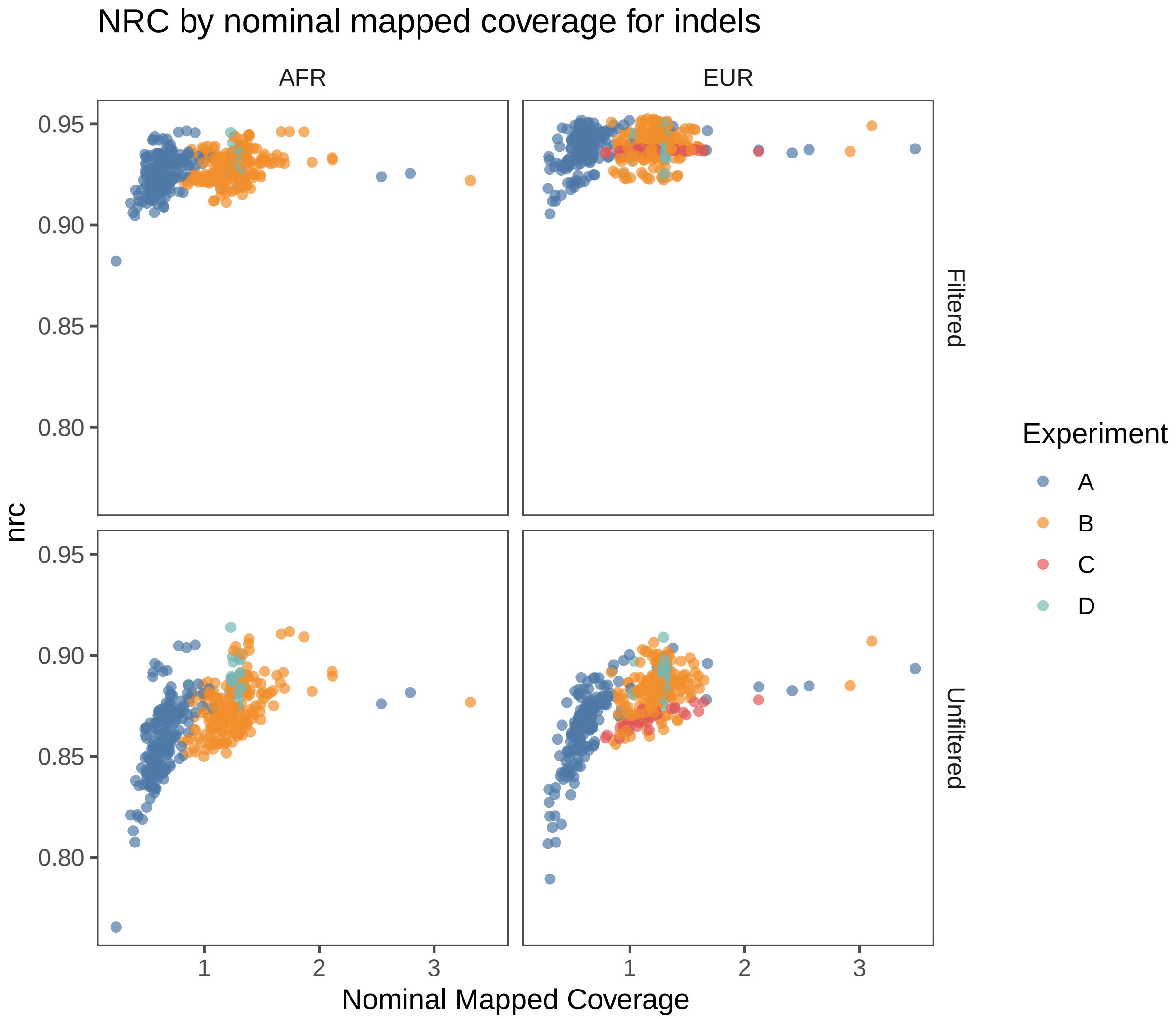


**Supplementary Figure 7:** Non-reference concordance for indels by nominal coverage for both filtered and unfiltered variant calls.

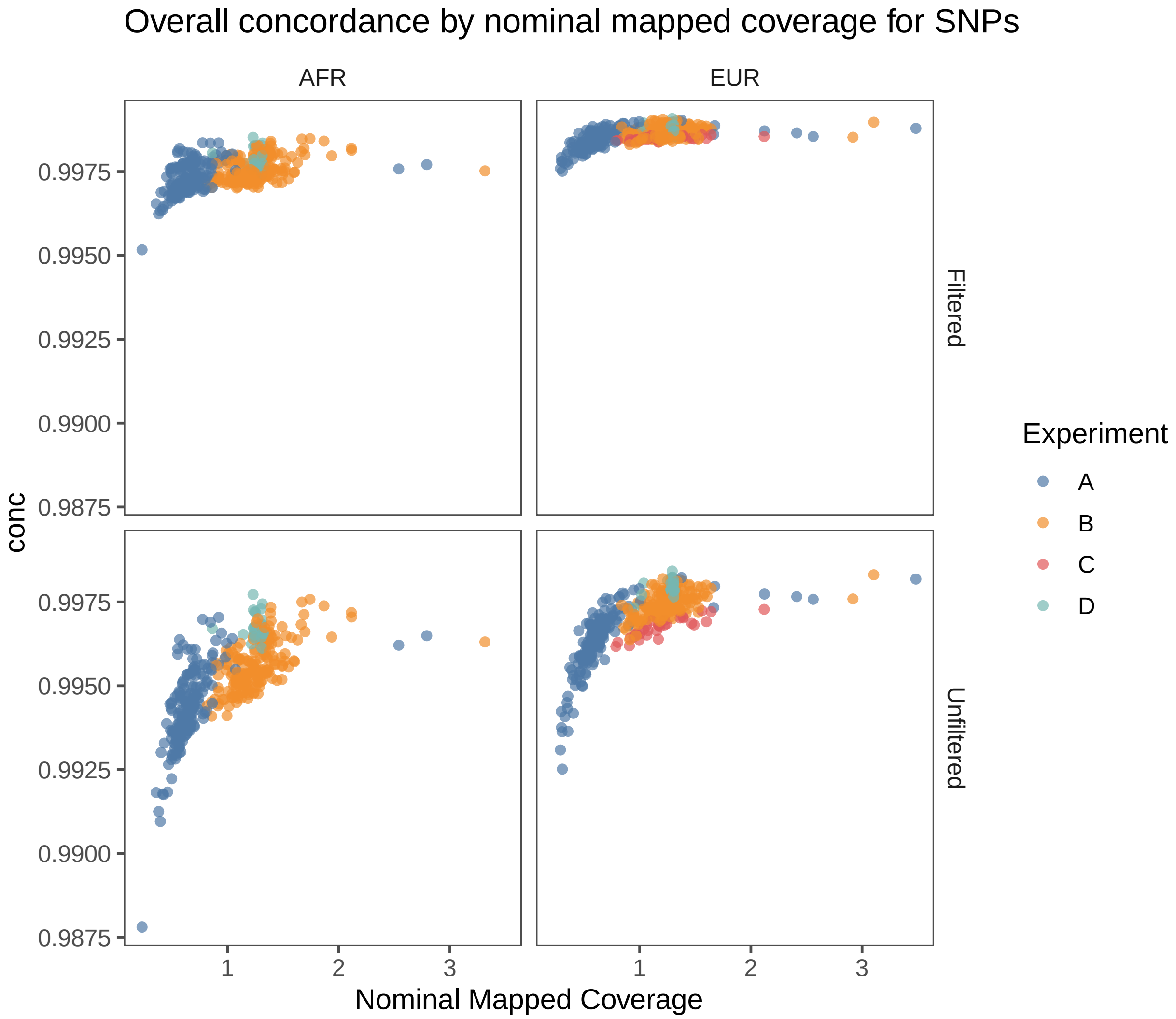


**Supplementary Figure 8:** Overall concordance for SNPs by nominal coverage for both filtered and unfiltered variant calls.

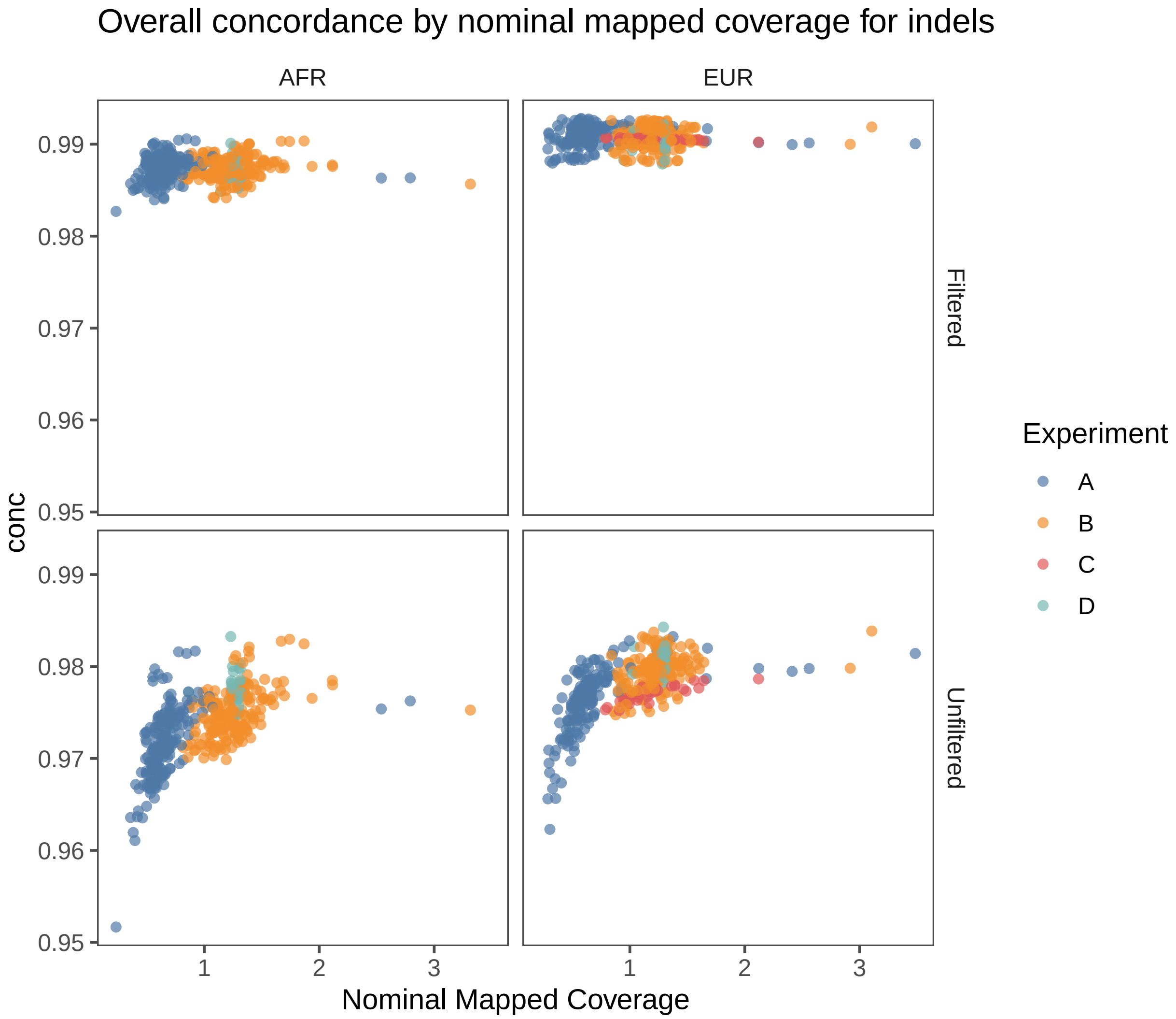


**Supplementary Figure 9:** Overall concordance for indels by nominal coverage for both filtered and unfiltered variant calls.

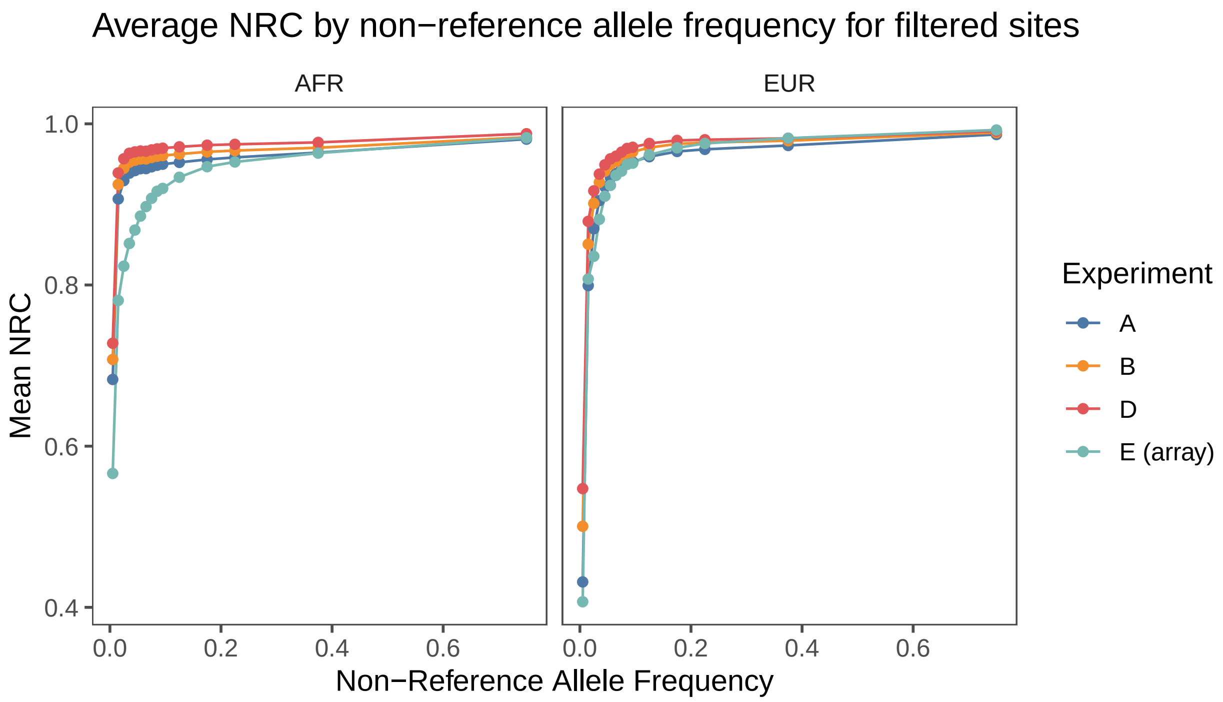


**Supplementary Figure 10:** Non-reference concordance for SNPs at filtered variant calls by non-reference allele frequency in the 1000 Genomes Phase 3 callset.

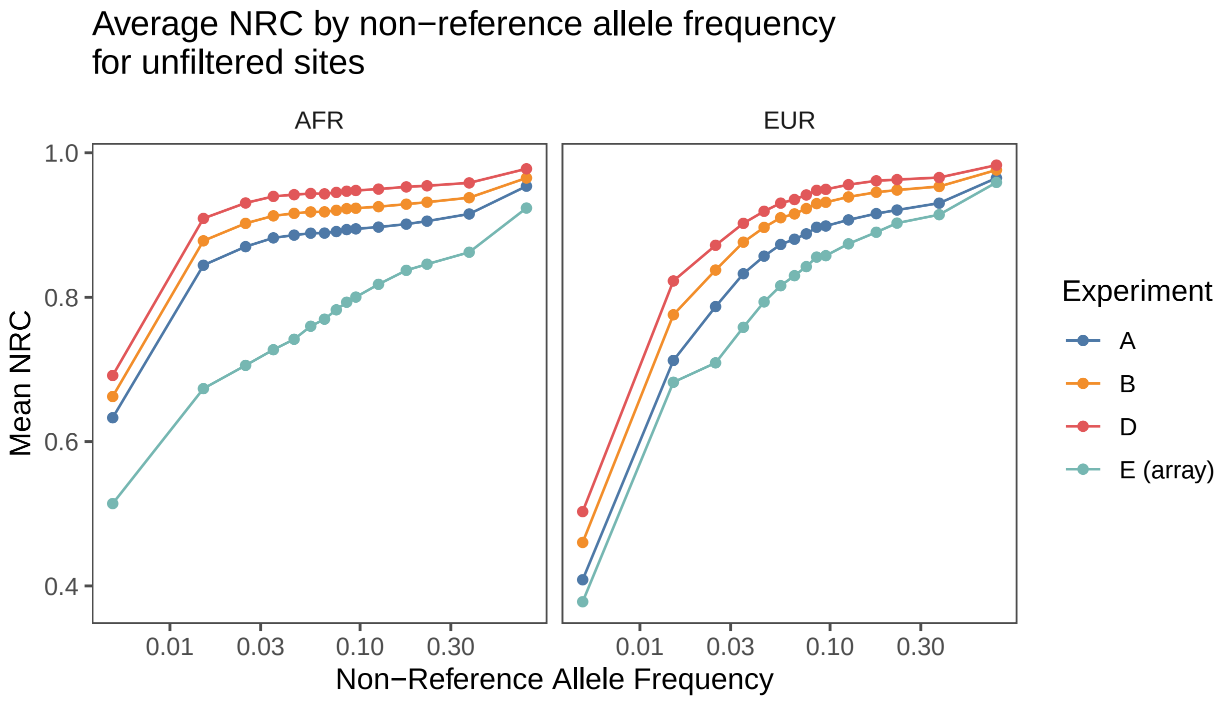


**Supplementary Figure 11:** Average non-reference concordance for unfiltered SNPs by superpopulation by non-reference allele frequency in 1KGP3. Same figure as Figure 2 on a log scale.

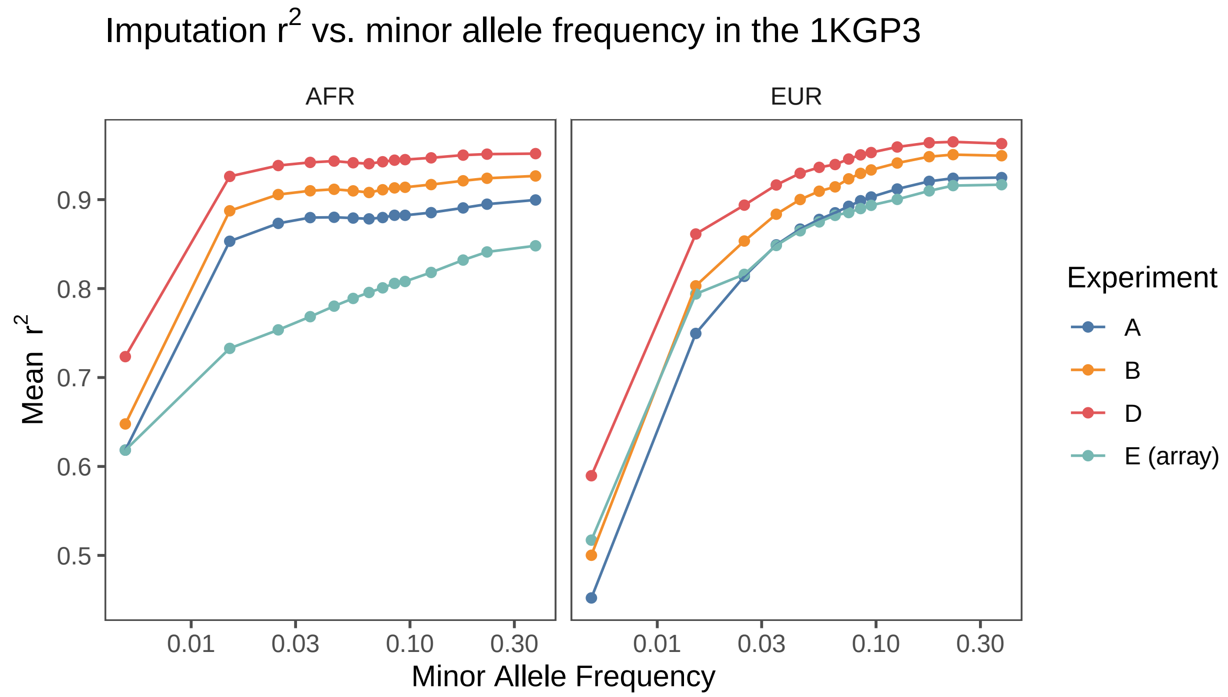


**Supplementary Figure 12:** Same as in Figure 3 but on a log scale. The values on the $x$-axis are the midpoints of the minor allele frequency bins. The bin edges are at 0, 0.01, 0.02, 0.03, 0.04, 0.05, 0.06, 0.07, 0.08, 0.09, 0.1, 0.15, 0.2, 0.25, 0.5.

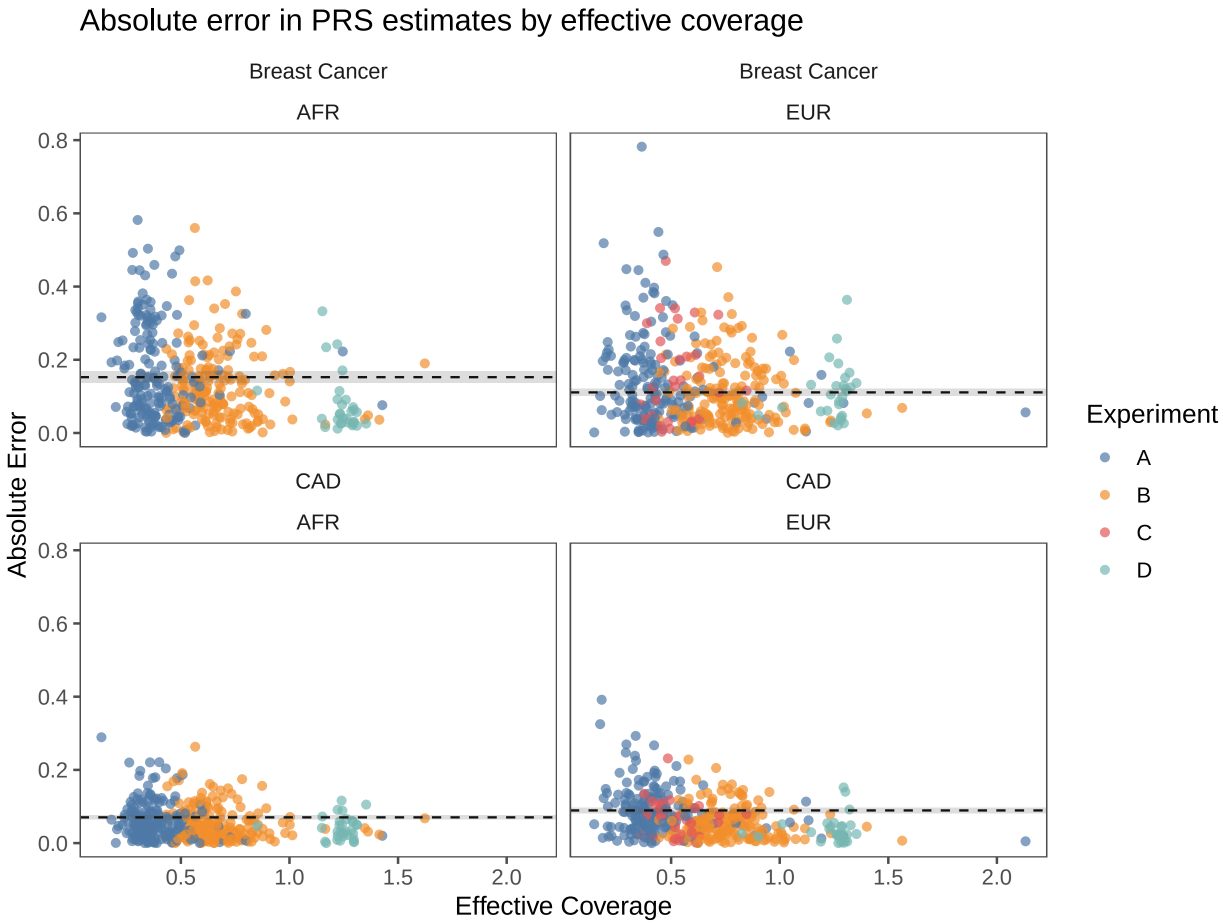


**Supplementary Figure 13:** Absolute errors in polygenic risk score estimates by effective coverage. The dotted horizontal lines denote the mean error in PRS estimation from Illumina GSA calls across cell lines replicates. The shaded area about each dotted horizontal line denotes the standard error of the mean of these quantities.

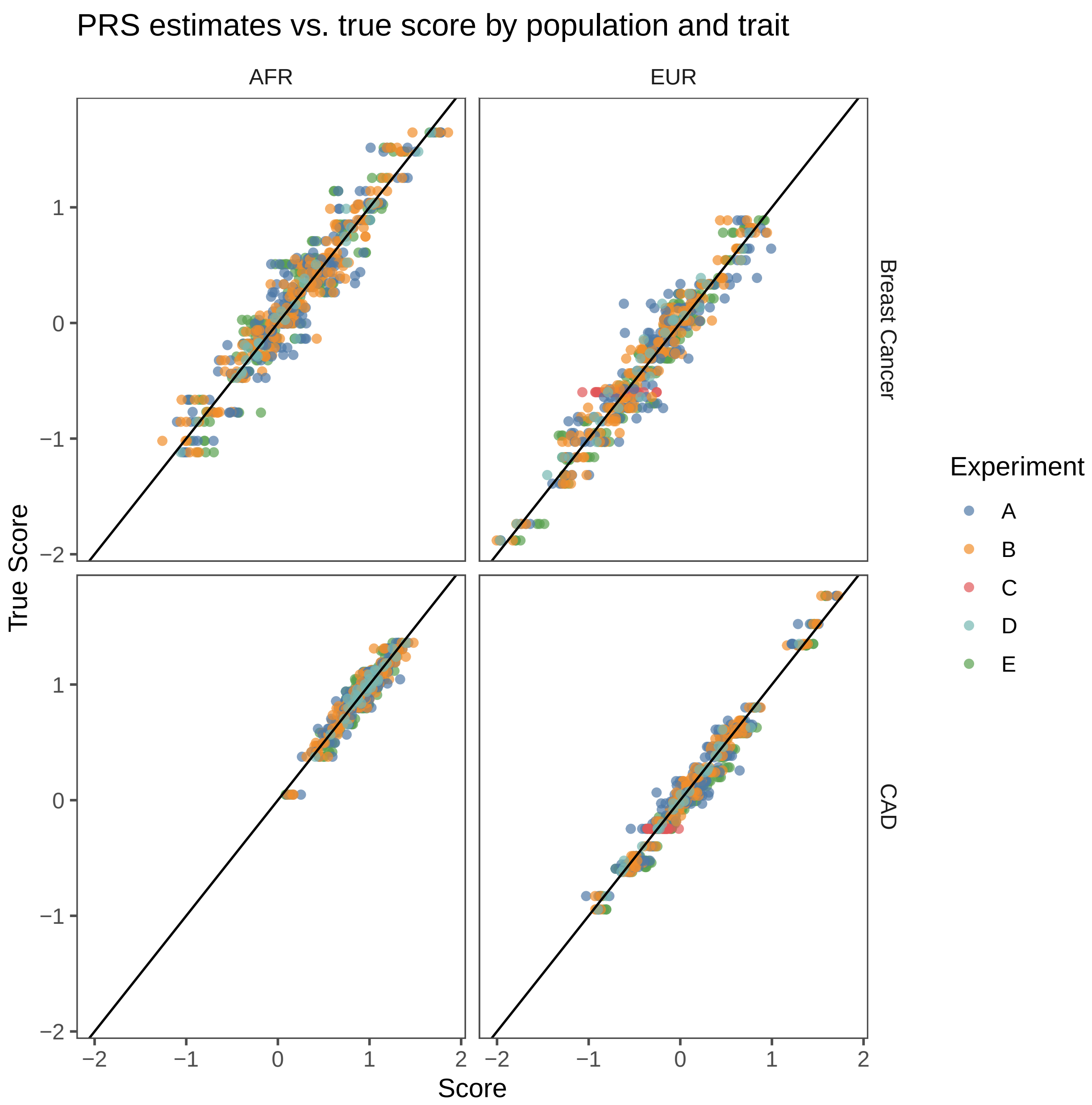


**Supplementary Figure 14:** Scatterplot of estimated PRS from imputed dosages vs the true PRS for all samples across experiments, populations, and traits. The PRS computed off genotypes in the 1KGP3 genotypes was considered the "truth" value for all cell lines.
